# Diatom Logic: Rewritable active nematic circuits in tidal-flat ecosystems

**DOI:** 10.64898/2026.09.14.751632

**Authors:** Qing Zhang, Manu Prakash

## Abstract

At the ocean-land interface lies a critical yet largely ignored ecosystem of tidal flats^1^. These habitats are colonized by diatoms that form thin surface biofilms, creating ecosystem-scale cellular collectives stretching meters to kilometers ^2–4^. How these collectives coordinate functions across vast spatiotemporal scales remains poorly understood. This task requires both long-distance transfer and long-term storage of information^5, 6^. In cellular collectives, trigger waves have been known to provide rapid, non-attenuating signaling^7–9^, while extracellular matrices allow cells to externalize structural memory^10–12^. Yet, how propagating signals and structural memory integrate to shape collective functions remains elusive. Combining field observations with laboratory reconstitution, we show that tidal-flat diatoms assemble active nematic circuits that physically wire trigger-wave routing into structural memory. Here, trigger waves recruit resting diatoms into polarized motion steered by the nematic director, and topological defects act as gates that focus or bifurcate propagating wave fronts. Memory follows a fast-read, slow-write rule: rapid signals deform the cellular nematic, while slowly evolving cell-deposited mucilage retains a hidden orientational template that guides alignment recovery. Strikingly, recurrent waves progressively remodel the nematic architecture, letting the collective reshape its future routing. Coupling rapid wave propagation to extracellular memory allows collectives to resolve the stability-plasticity dilemma, maintaining structural cohesion while adapting to recurrent stimuli. Our work reveals previously hidden adaptive nematic circuits that self-organize at ecological scales, establishing a physical framework for time-programmable active materials.

## Introduction

Tidal flats host dense microphytobenthic biofilms in globally extensive intertidal ecosystems^1,4,13^. These surface biofilms are often dominated by diatoms, single-celled algae that collectively span meters to kilometers across the landscape^2–4,14^. These diatoms contribute substantially to primary production, sediment stabilization, and biogeochemical exchange, functions that sustain coastal food webs and shoreline stability in environments repeatedly reworked by tides^15–17^. The dominant taxa within these surface layers are elongated pennate diatoms that glide over sediment while secreting adhesive extracellular mucilage^18–20^. While most mechanistic studies have focused on vertical migration, resuspension, and exchange with the overlying water^21–25^, far less is known about the architecture of this large-scale cellular collective as a densely packed surface film. Specifically, it remains unclear what principles govern the organization and how these cells coordinate collectively across the tidal-flat surface over many orders of magnitude in space and time. This missing link limits our ability to connect transient cell-scale mechanical behavior to ecosystem-scale biomass distribution and flux under rapidly fluctuating ecological conditions.

Coordinating the spatiotemporal collective functions at ecosystem scales requires both long-distance information transfer and long-term storage of past states. This necessity highlights two parallel pathways explored in cellular collectives: propagating signals and long-lived environmental states^5,6,10^. For information transfer, trigger waves provide a robust mechanism for long-range coordination because local positive feedback regenerates the front, allowing activation to propagate without progressive loss of speed or amplitude^8,9,26^. Trigger waves can be chemical, electrical, mechanical or hydrodynamic in nature^5,7,27–29^. Although the mechanisms that sustain these waves are increasingly well understood in lab model systems, how the physical architecture of the medium, particularly its anisotropy, shapes wave propagation remains underexplored^30–32^. Recent work shows that spatial heterogeneity^31,33^ and nematic cell alignment^32^ can organize and steer propagating fronts. However, it remains unclear whether trigger-wave passage can reciprocally remodel the macroscopic topological architecture that guides subsequent propagation. If such changes persist, this reciprocal coupling could encode communication history within a living collective.

For long-term information storage, one primary route for cellular collectives is externalized memory, in which past activity is encoded in persistent changes to the extracellular environment ^10,11,34^. These environmental traces can retain spatial or orientational information and feed back on subsequent collective organization^10–12,35^. In tidal-flat diatoms, secreted extracellular mucilage provides a natural candidate substrate for such externalized memory. While rapid trigger-wave communication and persistent environmental memory each have been recognized in biological information processing, how they integrate to shape collective function in a non-neuronal living material remains unresolved.

Here, we establish tidal-flat diatom mats as an ecosystem-derived active material that realizes this integration. Building on field observations of native biofilms, we demonstrate that diatoms self-organize into largely arrested, defect-rich nematic circuits that route trigger waves. These waves recruit resting cells into locally polarized gliding, with the macroscopic wavefront consistently outpacing individual cell motility. Unlike traditional systems, these trigger waves in diatoms carry both information and mass flux (motile diatoms). Remarkably, we uncover a fast–slow mechanism coupling two orientational fields: a rapidly deformable, excitable cellular nematic and a slowly evolving, cell-deposited anisotropic mucilage matrix. While wave propagation rapidly deforms the cellular nematic pattern, the mucilage matrix preserves a hidden orientational template for recovery. Recurrent waves progressively remodel the nematic circuit, physically rewriting the topology of routes available to later signals. This system reveals self-built topological trigger-wave circuits with externalized material memory, linking history-dependent information flow to the macroscopic organization of a fluctuating coastal biofilm.

## Results

### Ecosystem-derived, multiscale nematic mosaics

Through field studies with on-site microscopy, we establish that native tidal-flat diatoms in the microphytobenthos can form striking macroscopic architectures. To define these architectures, we examine intact surface layers from Aardvark Beach and Duck Pond in California, United States; and Masian Mudflat, in Korea (see Methods). To our surprise, across sites tens of meters apart, pennate diatoms align along their long axes without apparent periodic positional order, forming nematic textures (Fig. 1a). Similar textures recur across sites and seasons (Fig. S1), indicating that nematic organization is a reproducible feature of the native tidal-flat diatom communities. To determine whether this reproducible nematic organization is intrinsic, we isolated wild morphotypes, provisionally identified as *Cylindrotheca* sp. and *Navicula* sp., and established laboratory cultures from two sites in California. Both isolates reconstitute nematic mats on glass and plastic substrates, forming densely packed, locally aligned domains with no periodic positional order (Figs. 1b, S2-3). Tidal-flat diatoms thus form a living liquid-crystalline material both in the lab and field conditions. Scanning electron microscopy further shows nematic alignment of *Cylindrotheca* cells (Fig. 1c). Because *Cylindrotheca* sp. remains stable in long-term culture, subsequent analyses focus on this isolate.

**Figure 1.**
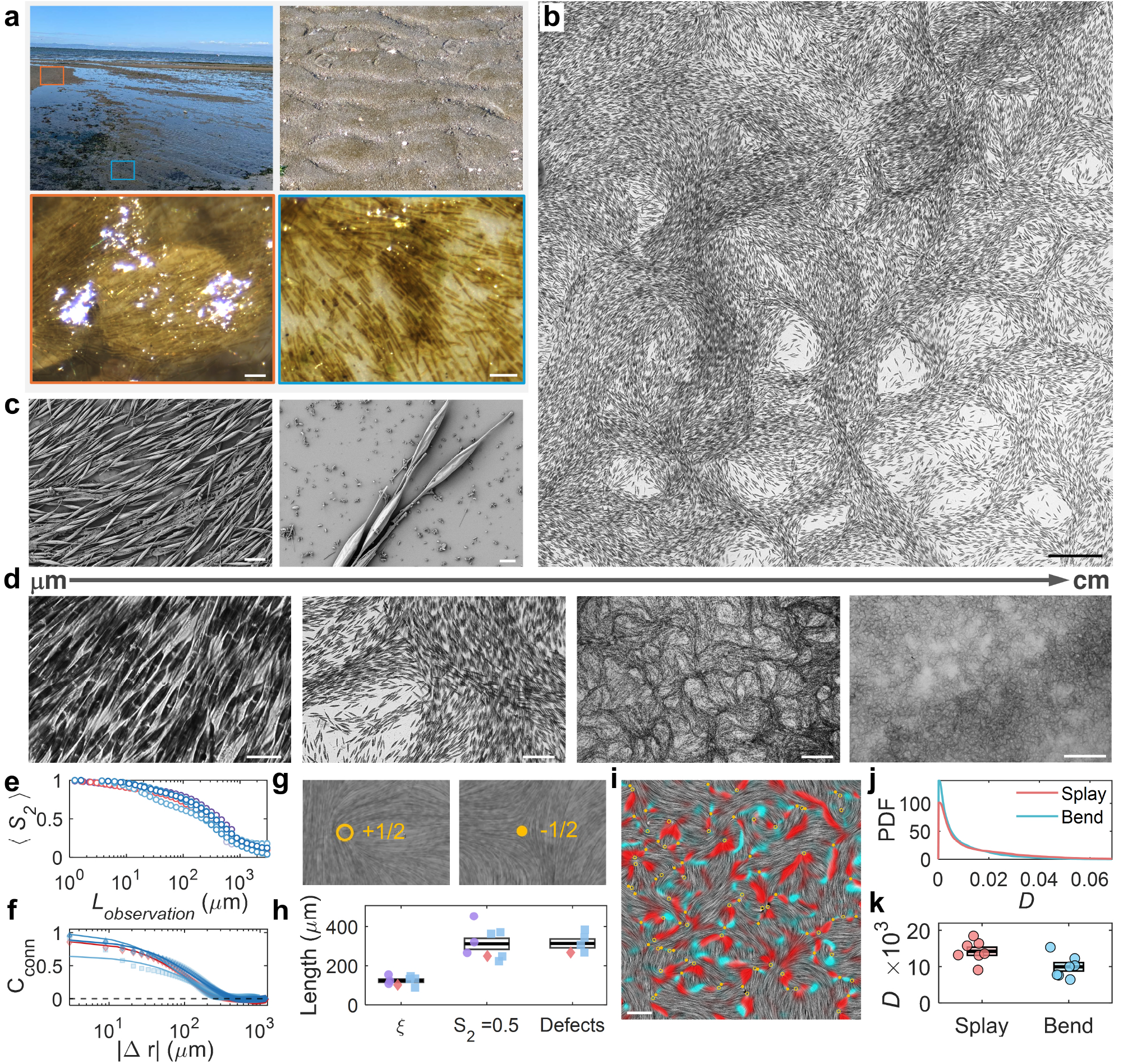
Multiscale nematic organization in tidal-flat diatoms. (**a**) Field observations (Aardvark Beach, CA). Upper left: two sampling sites ~ 30 m apart, marked by orange and blue boxes. Upper right: biota layer on tidal flat. Lower panels: microscopy of orange (left) and blue (right) sites shows locally aligned diatoms widespread across the ecosystem. Scale bars: 50 *µ*m. (**b**) Isolated *Cylindrotheca* sp. spontaneously forms nematic patterns. Scale bar: 200 *µ*m. (**c**) Scanning electron microscopy (SEM) reveals densely packed, parallel alignment (left) and fusiform cells with tapered ends and helical frustules (right). Scale bars: 20 *µ*m and 5 *µ*m. (**d**) Multiscale nematic organization from cell-scale alignment to centimeter-scale patterns. Scale bars from left to right: 20 *µ*m, 200 *µ*m, 1 mm, and 1 cm. (**e**) Nematic order *S*_2_p*L*q over observation windows *L* across magnifications (purple: 20×, red: stitched 20×, blue: 4×). (**f**) Connected orientational correlation *C*_conn_(*r*) versus radial separation Δ**r**; solid lines are exponential fits yielding correlation length *ξ*. (**g**) Representative ±1/2 defects in director fields visualized using line-integral convolution (LIC). (**h**) Comparison of *ξ*, half-order scale 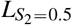, and defect-density length, *ℓ*_defect_, across individual 20× 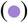, stitched 20× 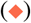 and 4× 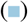. Shorter *ξ* reflects local orientational decorrelation; similar 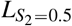 and *ℓ*_defect_ suggest a link between mesoscale loss of nematic order and defect organization (boxes: mean ± standard error of the mean (s.e.m.)). (**i**) Splay (red) and bend (cyan) deformations mapped on LIC director fields with defects (yellow). (**j**) Probability density functions (PDF) of dimensionless measures of splay and bend deformation, *D*. The heavier splay tail indicates more frequent large-splay deformations. (**k**) Higher mean *D* for splay than for bend indicates more pronounced converging and diverging distortions in the nematic pattern (boxes: mean ± s.e.m.).

Next we explore how this nematic order varies across multiple length scales (Fig. 1d; see Methods). Neighboring cells pack into aligned nematic patterns that curve, merge and diverge, assembling into a macroscopic mosaic of topological defects with curvilinear domains. To quantify this structural hierarchy, we measure the connected orientational correlation length (*ξ* = 123 ± 8 *µ*m, *n* = 8), the coarse-grained nematic order half-decay 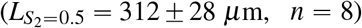, and the defect-density length of ±1/2 defects 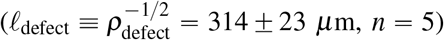 (Figs. 1e–h, S3, Table S1). The separation between the shorter local correlation *ξ* and the larger domain size 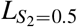 reflects strong local alignment embedded within a broadly curved texture. Additionally, 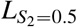 and *ℓ*_defect_ converge on the same characteristic length scale (~ 310 *µ*m). This agreement across observed magnifications links the mesoscale loss of nematic order to the spatial distribution of ±1/2 topological defects, suggesting that defects organize the characteristic scale of the nematic pattern (Fig. 1h).

To uncover the mechanics shaping these curvilinear domains, we decompose the director field into splay and bend deformations, which are strongly localized around topological defects and domain boundaries (Figs. 1i, S4; see Methods). Interestingly, splay deformations systematically dominate bend, exhibiting a broader high-value tail in distribution and a higher mean score (⟨*D*_splay_⟩ = (14.22±1.13) × 10^−3^ versus ⟨*D*_bend_ ⟩ = (9.95±1.16) × 10^−3^, *n* = 7; Fig. 1j,k). The highly tapered geometry of *Cylindrotheca* cells (Fig. 1c) provides a plausible microscopic contribution to this splay-rich architecture, potentially leading to the curvilinear nematic texture (see Discussion).

### Trigger waves in tidal-flat diatom nematics

Having established the nematic architecture of tidal-flat diatoms, we further investigate the dynamics of these living mosaics. Surprisingly, we uncover a previously unrecognized state in tidal-flat diatoms: spontaneous trigger waves that fluidize arrested nematic mats into collective motion (Movie S1). As visualized by arrival-time tracks, activation fronts nucleate locally and spread across the surrounding nematic mosaic (Fig. 2a). The activation produces pronounced cellular redistribution by mobilizing previously arrested cells within the diatom film. Eulerian velocity fields (**V**_*p*_) from particle image velocimetry (PIV) quantify the speed and direction of local population motion. These velocity fields reveal that, during wave passage, cells in neighboring regions near the wavefront move in locally coherent directions (Fig. 2b). Cell body axes remain locally nematically aligned during collective motion (Movie S1). Thus, the wave switches the mat from an arrested nematic state into locally velocity-polarized collective flow.

**Figure 2.**
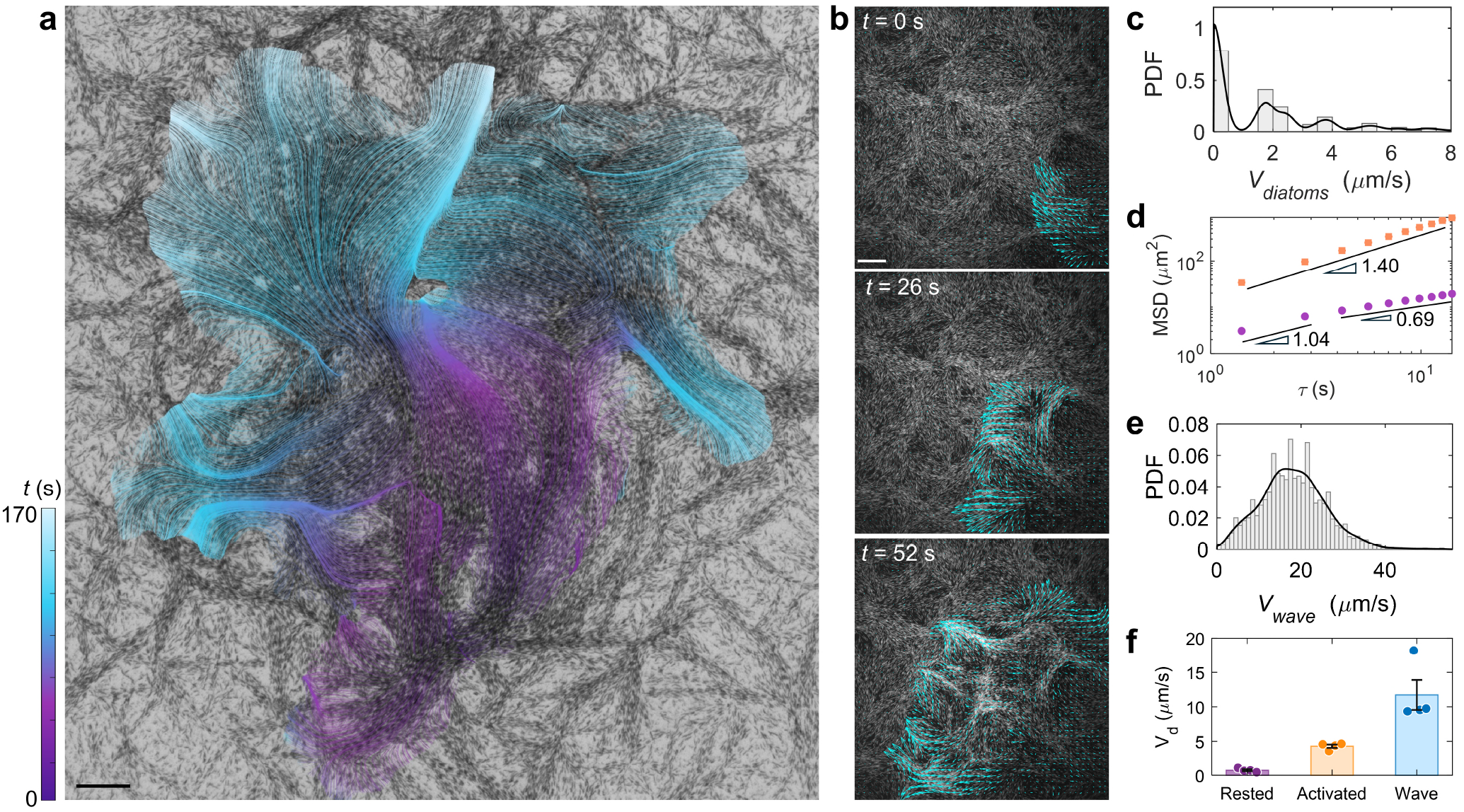
Trigger waves mobilize arrested nematic diatom collectives into locally velocity-polarized flow. (**a**) Spatiotem-poral propagation of a trigger wave through nematic tidal-flat diatoms (*Cylindrotheca* sp.). Tracks derived from population velocities are overlaid on a bright-field image. Color encodes local arrival time (purple to cyan), outlining a heterogeneous activated region. Scale: 200 *µ*m. (**b**) Eulerian velocity field from particle image velocimetry (cyan arrows), at times relative to the first frame. Near the propagating front, neighbouring vectors point in a common direction, indicating locally velocity-polarized collective flow. Scale: 200 *µ*m. (**c**) Probability density function (PDF) of individual gliding speeds, *v*_diatoms_. Grey bars are probability-density histograms and the black curve is a smoothed density estimate, revealing coexisting resting (near-zero speed) and activated motility (finite-speed peaks) states. (**d**) Mean-squared displacement, MSD (*τ*), for cells outside 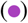 and inside 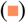 the wave. Symbols show mean ± s.e.m. across trajectories; error bars lie within the symbols. Power-law scaling, MSD ~ *τ*^*α*^, shows that the outside cells cross over from nearly diffusive motion (*α* ≃ 1.04) to subdiffusion (*α* ≃ 0.69), whereas the inside cells are superdiffusive (*α* ≃ 1.40). (**e**) Distribution of wave-front speeds for a representative trigger wave. (**f**) Comparison of resting-cell, activated-cell and mean wave-front speeds across independent experiments. Bars show mean ± s.e.m.; points denote biologically independent experiments (*n* = 4). The wave front travels much faster than individual gliding, consistent with a collective activation front rather than propagation by single-cell motility alone.

Next, we perform single-cell tracking, which reveals a sharp transition between motility states of individual diatoms (Movie S2; see Methods). Instantaneous cell speeds exhibit a nearly bimodal distribution, distinguishing cells trapped in the arrested state (near-zero speeds) from those released in the active gliding state (finite speeds) (Fig. 2c, Fig. S5). The nearly bimodal distribution supports distinct resting and activated motility states of diatoms. Mean-squared displacement (MSD) scaling (MSD ~ *τ*^*α*^) further resolves this transition (Fig. 2d). Cells outside the wave undergo short-time fluctuations (*α* = 1.04, nearly diffusive motion) but remain confined at the same position at longer lag times (*α* = 0.69, subdiffusion). In contrast, activated cells exhibit superdiffusive transport (*α* = 1.40), consistent with persistent gliding. Unique to our system, a trigger wave thus fluidizes the arrested nematic, transitioning cells from a stationary state to directionally persistent motion.

Strikingly, the activation front of the trigger wave outpaces individual cell motion. For the representative wave (Fig. 2b, Movie S2), local front speeds form a unimodal distribution centred near 18 *µ*ms^−1^ (Fig. 2e). Across four independent experiments, the mean front speed remains consistently high at 11.71±2.16 *µ*ms^−1^, which is nearly three times faster than the 4.30 ±0.24 *µ*ms^−1^ gliding speed of activated cells (Fig. 2f). This kinematic separation rules out transport of diatoms as a sole propagation mechanism, pointing instead to a self-propagating activation front (trigger wave) that outruns individual-cell motion.

To explain how the wavefront outruns individual cell motility, we provide a minimal decomposition of wavefront motion into directed transport and regenerative recruitment: 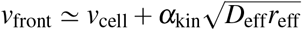, where *v*_cell_ is the activated-cell velocity, *D*_eff_ is the effective diffusivity of activation, 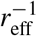 is the characteristic local recruitment time, and *α*_kin_ > 0 depends on nonlinear activation and recovery kinetics (see Supplementary Information). The scale 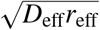 defines the speed at which spatial coupling recruits resting cells. Thus, while activated-cell motion provides directional transport, the activation field spreading ahead of the moving population sequentially recruits resting cells, allowing the macroscopic front to exceed individual-cell speeds.

### Nematic circuits route trigger-wave propagation

The trigger wave in tidal-flat diatoms does more than switch individual cells from rest to motion: it propels them through a structured transmission medium built by the cells themselves. Wave propagation through this curvilinear nematic mosaic is spatially heterogeneous (Fig. 2a, Movie S1). This heterogeneity suggests that the nematic director and local cell-density landscape jointly shape the accessible routes for collective activation. To test this idea, we co-register the near-front, wave-associated Eulerian population velocity field, **V**_*p*_, with the pre-wave nematic and density fields (see Methods). Regions of strong population motion preferentially coincide with aligned nematic bundles in dense parts of the mat (Fig. 3a,b).

**Figure 3.**
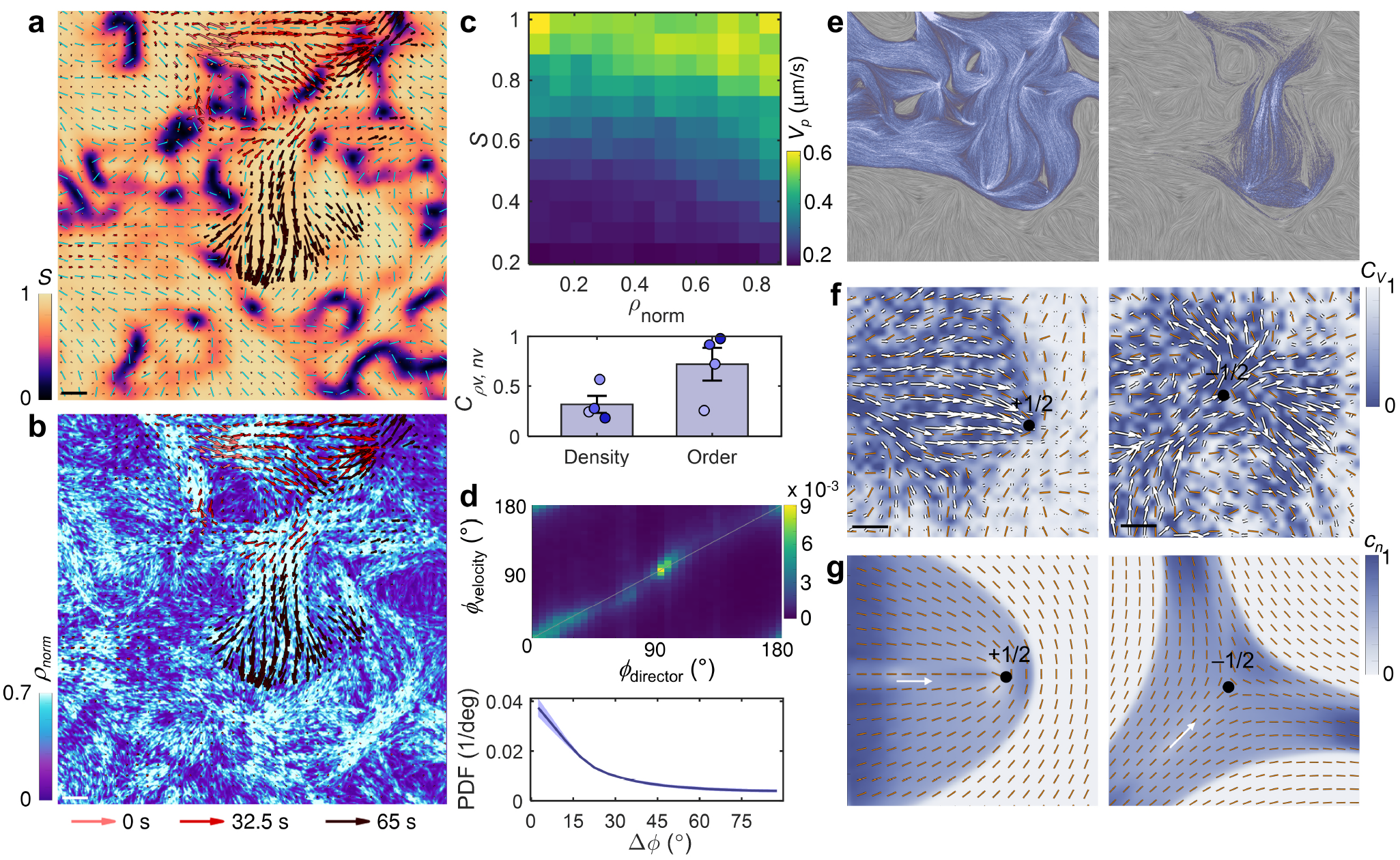
Nematic pattern and cell density shape trigger-wave propagation. (**a**,**b**) Local population velocity field, **V**_*p*_, measured by particle image velocimetry (PIV) during trigger-wave propagation in *Cylindrotheca* sp., overlaid on nematic order *S* (**a**; cyan rods, directors) and normalized density *ρ*_norm_ (**b**). Arrow colors mark *t* = 0, 32.5 and 65 s. Scales: 100 *µ*m. (**c**) Dependence of local speed, *V*_*p*_ = |**V**_*p*_|, on *ρ*_norm_ and *S*. Heat map: mean of per-experiment median *V*_*p*_ binned by *ρ*_norm_ and *S*. Bar plot: *V*_*p*_ correlates more strongly with nematic order (*C*_*nv*_) than density (*C*_*ρv*_); bars show the mean, error bars indicate s.e.m., and points denote independent datasets (*n* = 4). (**d**) Velocity orientation, *ϕ*_velocity_, preferentially follows director orientation, *ϕ*_director_: the joint distribution lies near the diagonal (upper) and the angular mismatch, Δ*ϕ* = arccos (|cos(*ϕ*_velocity_ − *ϕ*_director_)|), peaks near zero (lower). (**e**) Stochastic propagation model on the measured director and density fields. Left: nematic guidance only; right: nematic guidance with density gating. Grey: measured director field rendered by line-integral convolution; blue–white shading: cumulative simulated propagation footprint. Including density dependence better captures the experimental activated region. (**f**) Topological defects act as nematic lenses: the converging sector of a +1/2 defect focuses an incoming trigger-wave front, whereas the threefold −1/2 texture fans it outward. White arrows: **V**_*p*_; blue shading: activated region identified using the local PIV cross-correlation coefficient *C*_*v*_; orange rods: directors; black dots: defect cores. Scale bars, 100 *µ*m. (**g**) Simulations on idealized ±1/2 fields reproduce defect-mediated focusing and fan-out of trigger waves. Colormap: normalized cue field *c*_*n*_ = *c*/*c*_max_.

Next, we map the population speed, *V*_*p*_ = |**V**_*p*_|, against local nematic order (*S*) and normalized cell density (*ρ*_norm_). The two-dimensional map reveals a joint dependence: *V*_*p*_ increases most strongly with nematic order, while at low-to-intermediate *S*, higher density is associated with faster collective motion (Fig. 3c). Even in dense regions, *V*_*p*_ remains low at the lowest values of *S*, indicating that density alone is insufficient for strong coherent motion. Across four independent datasets, *V*_*p*_ correlates more strongly with nematic organization (*C*_*nv*_ = 0.72±0.16) than with density (*C*_*ρv*_ = 0.32±0.08). Additionally, we examine the relationship between the wave-associated population direction and the pre-wave nematic director. We compare the local velocity orientation, *ϕ*_velocity_, with the pre-wave nematic director orientation, *ϕ*_director_, using the axial mismatch Δ*ϕ* = arccos (|cos(*ϕ*_velocity_ − *ϕ*_director_)|). Their joint distribution lies tightly along the diagonal, and Δ*ϕ* peaks sharply near zero (Fig. 3d). The direction of wave-induced motion therefore follows the local director, indicating that the nematic mosaic operates as a self-assembled network of ‘nematic circuits’, guiding mass and information flow.

To further clarify the distinct roles of nematic alignment and cell density, we simulate wave propagation on the experimentally measured director and density fields (see Methods). With nematic guidance alone, the simulated footprint follows the director but extends beyond the experimentally observed wave-swept region. Introducing a density-dependent propagation probability suppresses propagation through low-density regions, producing a footprint that more closely matches the experimental wave-swept region (Fig. 3e, Fig. S6, Movie S3). Thus, the director guides propagation direction, while local density restricts which routes remain accessible.

The spatial organization of the director field around topological defects introduces an additional mesoscale control over wave propagation. Interestingly, in the nematic mosaic, topological defects do not merely mark static distortions; they reshape the geometry of information flow. To illustrate how defect geometry shapes wave routing, we examine representative encounters between wavefronts and ±1/2 defects (Fig. 3f, Movie S4). In the +1/2 defect example, wave-associated motion converges toward the defect core into a focused pathway. In the shown −1/2 geometry, the activated region splits and fans out along multiple directions. Simulations initialized with the corresponding defect geometries and wave-initiation sites reproduce these experimentally observed modes (Fig. 3g). Topological defects therefore act as biological ‘nematic lenses’, converting local topology into distinct patterns of wavefront focusing and spreading. Ultimately, biological information flow is routed through this defect-organized nematic circuit: director alignment steers the signal, density gates accessibility, and topological defects shape the wavefront geometry.

### Mucilage-encoded nematic memory

Performing long-term imaging, we find that trigger-wave activation in nematic tidal-flat diatom mats is not a single event; instead, waves recur and shape the cellular architecture (Movie S5). An initial wave propagates through the mat, redistributing cells along its path (Fig. 4a, Movie S5). Right after passage of a wave, both the normalized nematic order (*S*_*n*_) and the effective density-occupied area fraction ( *f*_*ρ,n*_) decrease within the wave-affected region (Fig. 4b), indicating reduced nematic alignment and biomass redistribution. Remarkably, ~8 h later, the cellular nematic mat reforms and is capable of supporting the propagation of a second wave along a similar pathway (Fig. 4a). Thus, the nematic architecture is transiently perturbed but not permanently erased. Such a living nematic circuit shows repeated cycles of deformation and recovery.

**Figure 4.**
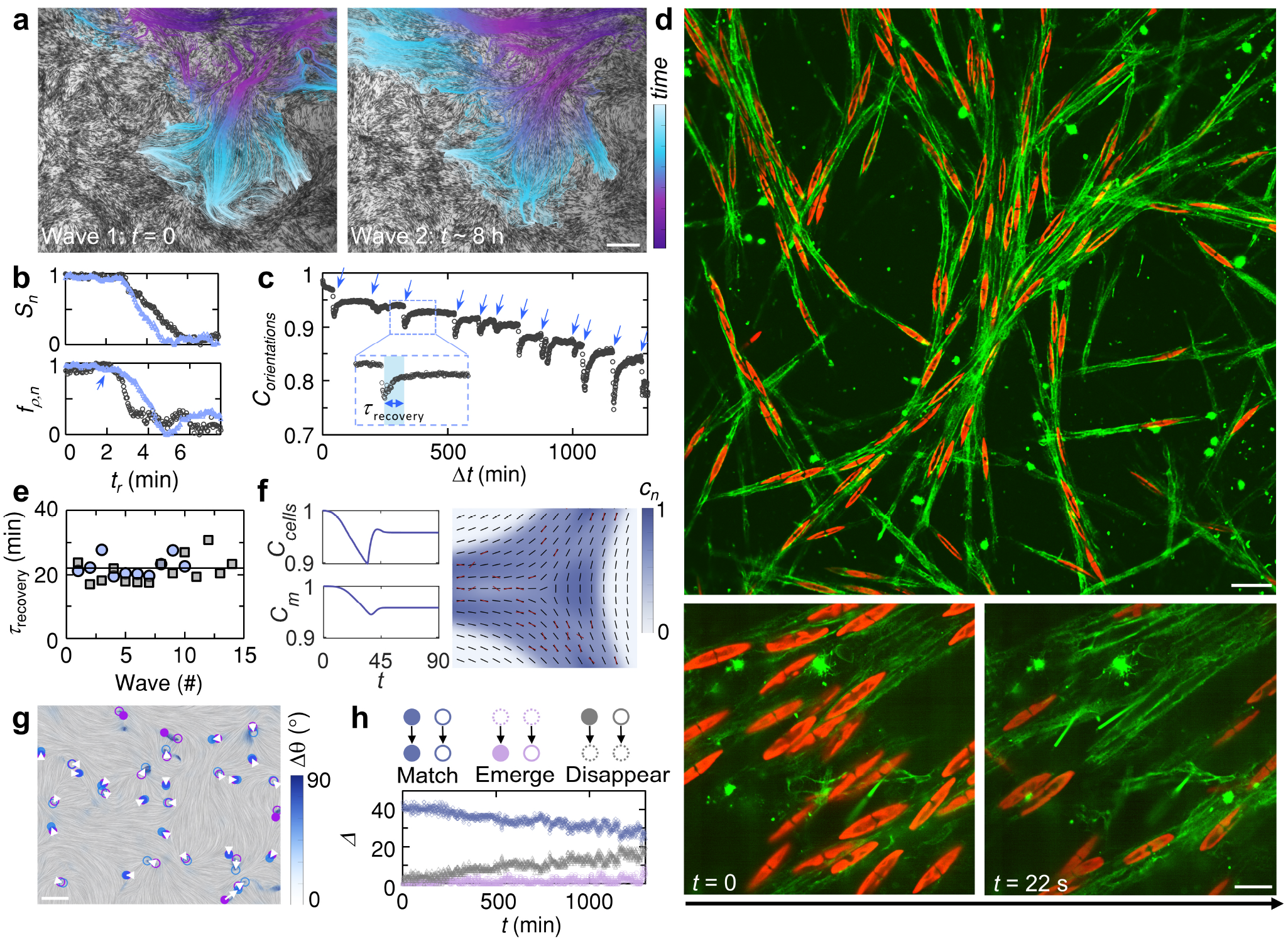
Mucilage-encoded memory supports recovery and remodeling. (**a**) Recurrent waves in the same field, ~8 h apart; color encodes local arrival time. Both waves follow similar propagation pathways. Scale: 200 *µ*m. (**b**) Normalized nematic order, *S*_n_ (upper), and effective density-occupied area fraction, *f*_*ρ*,n_ (lower), decrease during both wave passages in (**a**). Arrow: trigger point. (**c**) Cross-correlation of nematic orientation field, *C*_orientations_, drops abruptly at waves (blue arrows) and subsequently recovers. Inset defines the recovery time, *τ*_recovery_ from post-wave minimum to plateau. (**d**) Wheat germ agglutinin (WGA) staining shows oriented mucilage (green) deposited by diatoms (red). Lower: During a wave, cells move away while mucilage retains its orientation. Scales: 20 *µ*m (upper), 10 *µ*m (lower). (**e**) Recovery times, *τ*_recovery_, are nearly constant for successive trigger waves in two biological replicates 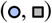 and comparable between them, consistent with the model prediction; black line, constant fit. (**f**) Coupled cell–mucilage simulations capture memory and remodeling: cross-correlation of cell orientation field, *C*_cells_, drops and partially recovers, enabled by slowly changing mucilage matrix (cross-correlation, *C*_*m*_), whose small persistent loss leaves residual remodeling. Right: pre-wave directors (black), mismatched post-wave directors (red). *t*: simulation time. Colormap: normalized cue field, *c*_n_ = *c/c*_max_. (**g**) Director-field remodeling between recurrent waves. Color map shows axis-folded mismatch, Δ*θ*, between pre-wave states of waves 1 and 2, overlaid on wave-1 directors rendered by line-integral convolution (LIC). Wave-1/2 defects are purple/blue; open/filled symbols mark +1/2/−1/2 defects; white arrows link matched defects. Scale: 200 *µ*m. (**h**) Defect fates during repeated trigger waves. Defect counts, Δ, by category relative to the first analyzed frame: matched defects decrease, disappeared defects increase, and emerged defects remain sparse, indicating progressive remodeling of the defect landscape.

To quantify the recovery, we measure the temporal cross-correlation of the nematic orientation field relative to the pre-wave pattern, *C*_orientations_ (*t*), which captures local director axis preservation (Fig. 4c). Each trigger wave (blue arrows) produces an abrupt drop in *C*_orientations_, reflecting rapid cellular nematic field deformation. Post-wave, the correlation recovers to a plateau below its pre-wave baseline. This partial recovery demonstrates a structural memory of the nematic architecture, but not exact restoration: each wave reads an existing orientational template, yet leaves behind residual structural changes permanently in the nematic circuit.

How does a microbial collective recover its previous nematic pattern after cells move away and local alignments are largely disrupted? One natural candidate is the extracellular mucilage secreted during raphe-associated diatom gliding, which mediates cell–surface interactions^19,36,37^. In our *Cylindrotheca* sp. isolate, SEM reveals a longitudinal raphe and abundant extracellular material coating and bridging cells, consistent with a raphe-associated mucilage system (Figs. 1c, S7). As cells move, they deposit polymeric mucilage trails that persist, potentially recording previous orientations. To test this, we stain the nematic diatom mat with wheat germ agglutinin (WGA) to label glycoconjugate-rich extracellular material (see Methods). WGA staining clearly reveals an oriented extracellular matrix matching the cellular nematic architecture (Fig. 4d). During wave passage, cells abruptly relocate, but the mucilage matrix largely remains and preserves previous orientation (Fig. 4d and Movie S6). Thus, while wave-induced cellular redistribution transiently disrupts the cellular director field (*ϕ*_*c*_) behind the advancing front, the stationary mucilage director (*ϕ*_*m*_) retains a hidden orientational template. The resulting orientational mismatch provides a mechanism for cellular realignment and governs the kinetics of recovery.

Defining recovery time, *τ*_recovery_, from the post-wave *C*_orientations_ minimum to its plateau (Fig. 4c, inset), we find *τ*_recovery_ remains nearly constant across successive waves and is comparable between independent experiments (Fig. 4e). We rationalize this reproducible timescale using a local angular-relaxation model adapted from a hydrodynamic theory of an active nematic coupled to a remodelable matrix^12^. The local cell–mucilage angular mismatch, *δ* = *ϕ*_*c*_ − *ϕ*_*m*_, evolves as 2∂_*t*_*δ* = − (*λ*_pin_+*λ*_remodel_) sin 2*δ*. Here, *λ*_pin_ = *kβ*_*m*_*ρ*_*m*_/(*β*_*m*_*ρ*_*m*_ +*β*_*c*_*ρ*_*c*_) drives cellular realignment toward the mucilage template, and *λ*_remodel_ = *k*_−_*ρ*_*c*_ captures slow rewriting of the mucilage template by cells. *β*_*c*_ and *β*_*m*_ are the effective cell–cell and cell–matrix alignment strengths, respectively. *ρ*_*c,m*_ are the cellular and mucilage matrix densities. *k* is the cellular reorientation rate, and *k*_−_ controls matrix rewriting. The model gives a characteristic angular-relaxation time *τ*_*δ*_ = (*λ*_pin_ + *λ*_remodel_)^−1^ (see Supplementary Information). In the matrix-pinned limit during post-wave recovery, *λ*_pin_ ≫ *λ*_remodel_, so 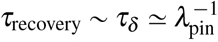. For a fixed effective pinning rate, the model predicts a constant recovery timescale, in agreement with experimental measurements (Fig. 4e).

Notably, recovery is just part of the story. During trigger-wave passage, activated cells rearrange relative to the persistent mucilage template. The resulting cell–matrix orientational mismatch can drive slow matrix rewriting, allowing structural plasticity to accumulate over recurrent waves and progressively remodel the nematic circuit. Coupled cell–mucilage numerical simulations under excitable waves capture this mechanism (Fig. 4f; see Methods). Activation induces nematic-guided motion, sharply dropping the cross-correlation of cell-orientation field, *C*_cells_. Conversely, the cross-correlation of mucilage-orientation field, *C*_*m*_, remains high as the slowly evolving matrix preserves the pre-wave template. Post-wave, this template drives the partial recovery of *C*_cells_. The small residual loss in *C*_*m*_ prevents complete recovery, consistent with gradual remodeling of the memory field.

Spatially, the remodeling of the nematic architecture is highly localized. Comparing relaxed director fields immediately before representative waves 1 and 2 (Fig. 4a) reveals that large orientation changes (Δ*θ*) concentrate around topological defects (Fig. 4g, S8; see Discussion). Time-resolved defect tracking further resolves the dynamics of the residual remodeling: there is a long-term increase in defect displacement despite partial reversals after individual waves (Fig. S9, Movie S4).

Classifying defect fates over repeated waves shows a decrease in matched defects, an accumulation of disappeared defects and sparse emergence (Fig. 4h). These changes suggest progressive simplification of the defect-rich nematic architecture (see Discussion). Because defects lens trigger waves, this slow turnover updates the routing landscape for future activation fronts. In the upper region, for example, defect loss coincides with a smoother director pattern, and the second wave expands where the first wave stalled (Fig. 4a).

Ultimately, the tidal-flat diatom mat operates as an adaptive memory of past events. Waves drive rapid cellular information transfer and dispersal of diatoms across a quasi-two-dimensional field, while the slowly deforming mucilage retains the structural memory. Because the mucilage orientation persists, the original activation pathways are largely retained in the short term. Over repeated cycles, however, the wave passage evolves due to topological operations, modifying the matrix and its defect landscape. The diatom collective system thus behaves as an excitable topological circuit where material organization guides information propagation, yet that same organization is progressively modified by the passage of past trigger waves.

### Programming excitation and nematic architecture

The adaptive diatom mats discovered here rely on two separable ingredients: a locally excitable trigger and a structural nematic routing field. To probe wave initiation, we applied three distinct perturbations: mechanical, chemical, and optical, each of which can nucleate an activation front (Fig. 5a, Movie S7). First, gentle indentation with a micropipette triggers activation from the point of contact. Second, localized injection of *trans,trans*-2,4-decadienal initiates collective motion. While this stress-signaling aldehyde is known to alter mucilage-mediated adhesion and traction^38,39^, we use it here as an experimental perturbation rather than as evidence for the natural endogenous cue. Third, optical laser spot illumination also nucleates a wave, albeit via an unresolved photothermal or photochemical mechanism. In all three scenarios, collective motion rapidly escapes the stimulated site. The convergence of these diverse inputs highlights that excitability is a robust, intrinsic feature of the diatom mat.

**Figure 5.**
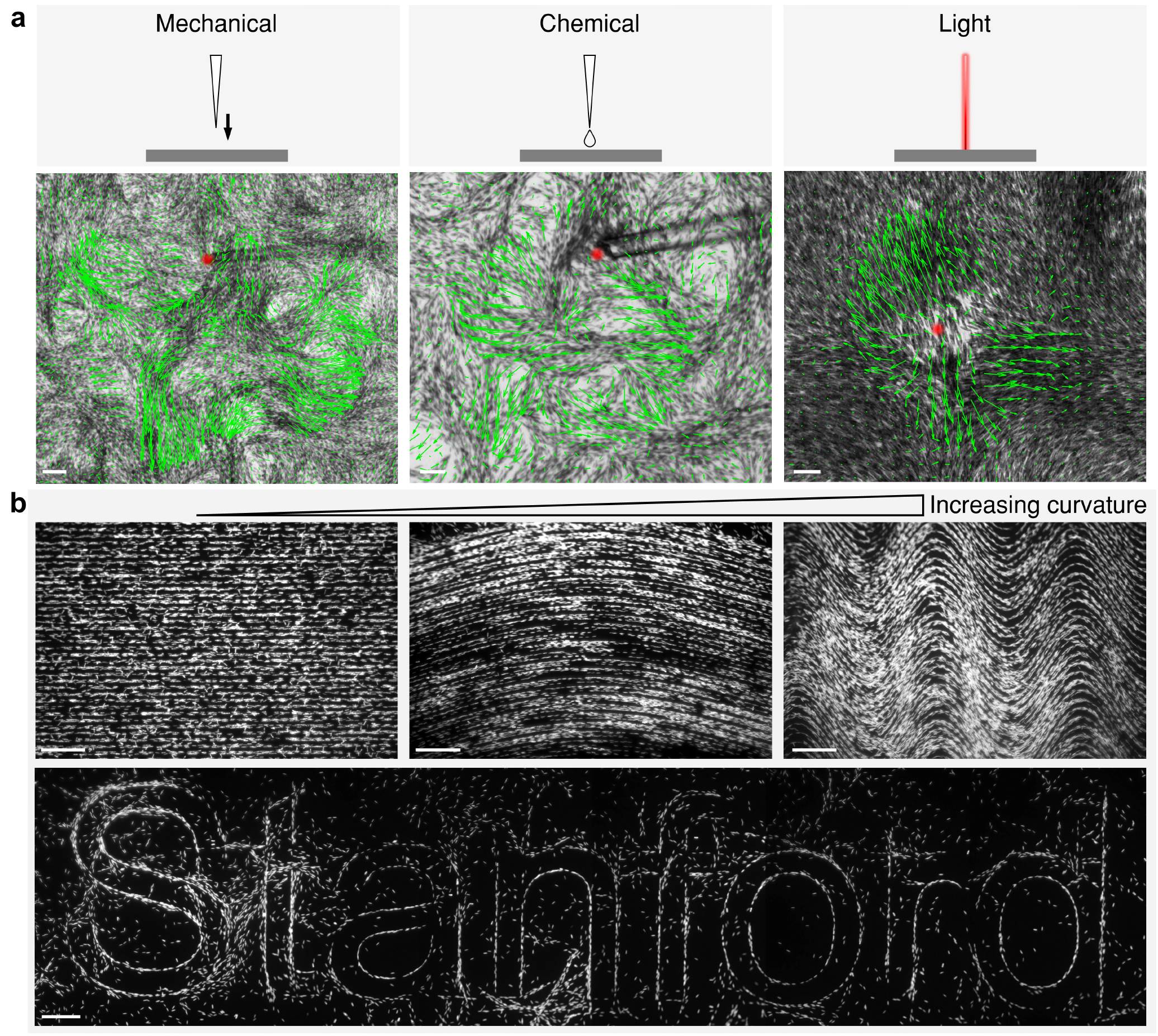
External control of wave initiation and alignment. (**a**) Trigger waves in nematically aligned *Cylindrotheca* sp. induced by local mechanical perturbation (left), chemical stimulation by *trans,trans*-2,4-decadienal injection (middle), and optical stimulation by laser illumination (right). Upper: Schematics show the applied perturbations. Lower: Experimental images show the resulting local population velocity field, **V**_*p*_, overlaid on bright-field images. Red marks the stimulation site and green arrows show **V**_*p*_, demonstrating that distinct local inputs can nucleate collective trigger-wave propagation through the surrounding nematic field. Scale bars: 50 *µ*m. (**b**) Designed surface grooves align tidal-flat diatoms into prescribed nematic architectures, providing topographic control of the collective orientation field. Straight grooves and sinusoidal grooves of different wavelengths (top), together with character-like topography (bottom), illustrate programmable diatom alignment and potential trigger-wave pathways. Scale bars: 200 *µ*m.

We next show that the routing medium can also be externally prescribed through initial conditions. On microgrooved substrates (average depth: 11–12 *µ*m), the typically self-assembled nematic mosaic organizes into a defined, engineered director field (Fig. 5b, S10). Cells align tangentially along straight, curved, and sinusoidal grooves (Fig. 5b, upper; S10a), and conform to curvature or character-like topography (Fig. 5b, lower; S10b). Surface geometry is thereby translated into a customizable living nematic architecture.

These two interventions, taken together, provide two complementary control axes over wave initiation and prescribed topological network, making designed wave routing through prescribed director fields feasible.

## Discussion and Outlook

Our results establish dense tidal-flat diatoms as a new ecosystem-derived nematic circuit where excitation, topology, and memory are functionally inseparable. The tidal-flat diatom mats form a splay-rich, slowly evolving defect mosaic in a frictional, history-dependent active-nematic regime that differs from other canonical active nematics^40^. The cells are substrate-bound, intermittently motile and coupled to a self-deposited extracellular matrix. The coexistence of a mobile cellular nematic and a slowly remodeling mucilage template gives the mat both fluid-like and solid-like features. The system may therefore represent an intermittently fluidized, matrix-coupled active nematic material. Confinement, substrate friction, gelation and matrix remodeling can stabilize ordered, jammed or frozen active-nematic states^11,12,41–43^. Excluded-volume interactions provide a baseline mechanism for nematic ordering of highly elongated rigid cells^44,45^, and contribute to shape-dependent alignment in active rods^46^, whereas tapered cell shape and mucilage-mediated adhesion help build the multiscale mosaic.

Within this nematic architecture, the trigger wave acts as both a biological signal and a motility-state transition, converting a largely arrested nematic into transiently velocity-polarized collective flow. Interestingly, while the structural order remains apolar, activation selects one of the two equivalent director branches and recruits collective motion along it. Anisotropic excitable-media models route wavefronts through prescribed anisotropic transport or orientation-dependent coupling^32,47^. In contrast, mechanochemical extracellular signal-regulated kinase (ERK) waves orient epithelial migration and cyclic adenosine monophosphate (cAMP) waves guide amoeboid chemotaxis, with net cell migration in both systems opposite to wave propagation^48,49^. The diatom wave combines and departs from both paradigms: a pre-existing apolar nematic guides the front, while the cells forming this propagation medium are themselves recruited into collective gliding in the direction of local front advance. The wave therefore mobilizes the same cellular architecture that guides it, generating locally polar mass transport that redistributes cell density and can contribute advectively to wavefront advance.

What switches the diatom collective from an arrested to a motile state in natural conditions remains an open question. In our diatom mats, mechanical contact, decadienal injection, and localized illumination each successfully nucleate a trigger wave. Prior literature links decadienal and nitric oxide to diatom stress responses and motility^38,39,50,51^. Because diatom gliding relies on raphe-associated adhesion^19,36^, a plausible working model posits that local stress rapidly alters cell–surface interactions, while a diffusible or mechanochemical cue recruits adjacent cells to switch to a motile state. The observation of finite propagation and wave stalling strongly suggests excitability thresholds governed by cell density or path geometry. Future simultaneous measurements of Ca^2+^ and NO dynamics, coupled with traction force microscopy, will disentangle these sub-cellular scale variables.

At the mesoscale, our experimental measurements and theoretical models provide a compact routing rule: the pre-wave director steers motion, local density gates route accessibility and defect geometry can focus or disperse the activation front. Topological defects thus act as ‘nematic lenses’. Yet they are not passive lenses: the largest persistent orientational changes between relaxed pre-wave states are concentrated near defects, while recurrent waves progressively modify the defect landscape (Fig. 4g,h). The accompanying reduction in nematic alignment (Fig. 4b) suggests a parallel with active nematic solids, where local melting of nematic order enables defect motion^52^. Extending our coupled cell–mucilage model to resolve interacting defects will clarify how recurrent excitation and coupling between the cellular and mucilage-matrix nematic fields reshape the defect landscape. Defects therefore both route the propagating signal and mark where its passage most strongly rewrites the routing landscape. This dual role extends their established functions in active flow, stress localization, cell fate and morphogenesis^53–58^ to a reciprocal coupling between biological information flow and topological remodeling.

Strikingly, the diatom circuits exhibit a fast–slow hierarchy: trigger waves rapidly mobilize cells and deform the cellular nematic, whereas the mucilage-coupled architecture evolves more slowly, guiding recovery and retaining residual structural changes. This kind of fast–slow separation is a recurring motif in neural and physical learning, in which rapid responses are shaped by slower variables that retain experience and tune future behavior^59–61^. Here, the persistent yet rewritable routing architecture positions the diatom mat as an ecosystem-derived platform for testing physical learning. Under controlled recurrent stimulation, wave passages would serve as training events, the slowly remodeled architecture as the learning degrees of freedom, and later wave trajectories as the readout. Independent control of wave initiation and nematic architecture provides the experimental handles to prescribe both stimulation history and routing geometry. Toward establishing the diatom mat as a physical learning system, future work will investigate whether prior wave traffic produces a retained improvement or specialization in later routing and biomass redistribution.

Ecologically, these nematic circuits provide a self-regulated, history-dependent mechanism for shaping biomass distribution in microphytobenthic systems. Trigger waves rapidly reconfigure the lateral footprint of surface biomass, while the persistent mucilage architecture preserves a template for recovery. A testable implication is a switch between mobilization and retention: if activation transiently weakens cell–surface adhesion, waves could prime cells for tidal entrainment and redeposition, whereas persistent mucilage could support biofilm cohesion and sediment stability between events^17,18^. Such fast–slow organization could convert past collective activity into a material legacy that biases future biomass retention and export, linking collective communication to ecological memory over large length scales.

Our findings identify a previously unrecognized mode of organization and coordination in the tidal-flat diatom mats examined here: ecosystem-derived nematic circuits built from cells and their extracellular matrix. Rapid signaling and persistent memory are often coupled within intracellular molecular networks at the cellular scale^62,63^. Here, this coupling is instead realized at the scale of the collective through a reciprocal signal–architecture loop, in which propagating signals and their guiding material pathways are co-constructed. The fast-read, slow-write dynamics of this loop offer a physical route to reconciling structural stability with adaptive plasticity in a non-neural living collective. This raises the prospect that an excitable living material could learn how to route its own signals at ecologically relevant length scales. More broadly, the system illustrates a distinctive paradigm for adaptive biological communication: an unwired cellular collective can build, store and revise its own information highways.

## Materials and Methods

### Field sampling, isolation, and cultivation of tidal-flat diatoms

We conduct field studies across multiple seasons at three sites: Aardvark Beach (San Mateo, CA) and Duck Pond (Palo Alto, CA) in the United States (April 2024; February, March and September 2025), and Masian Mudflat in South Korea (May 2024). During sampling, we excavate the top tidal-flat layer (approximately 10 10 cm in area and 2 cm in depth) to preserve the structural integrity of the surface biofilms. These intact samples are directly observed and recorded under a stereomicroscope (Zeiss) within 1–2 hours of collection to capture the native behaviors of the surface microbiome (see Supplementary Fig. S1 for detailed site images).

From the field samples collected at Aardvark Beach (37.59° N, 122.33° W), we isolate wild strains of benthic diatoms, specifically focusing on *Cylindrotheca* sp. We initially isolate approximately 150 individual cells, sequentially subculturing them as the populations expand to establish stable, robust cultures. These isolates are then cultivated in standard f/2 medium prepared with filtered seawater, and maintained in controlled environmental chambers at 20 °C under a 12-hour light/12-hour dark cycle.

### Sample preparation and imaging

To prepare samples for scanning electron microscopy (SEM), we fix the tidal-flat diatoms in a solution of 2.5% glutaraldehyde prepared in 0.1 M sodium cacodylate buffer (pH 7.2). Following fixation, we wash the samples three times in the same buffer to remove excess fixative, and then dehydrate them through a graded ethanol series (35%, 50%, 70%, 80%, 95%, and 100% v/v). Finally, the samples are critical-point dried for 60 minutes (Tousimis Research Corporation) and sputter-coated with a 3 nm layer of gold-palladium (60:40 ratio; Safematic) prior to imaging.

To visualize the underlying mucilage matrix, we fluorescently label the secreted mucilage. We first culture the diatoms for 1–2 weeks until a mature nematic biofilm develops. The mucilage is then stained by adding wheat germ agglutinin (WGA) conjugated to Alexa Fluor 555 (Thermo Fisher Scientific) at a 1:200 dilution, followed by a 2-hour incubation in the dark at 20 °C^64,65^. We select this fluorophore to minimize spectral interference from diatom autofluorescence.

We perform fluorescence imaging using a Nikon Eclipse Ti2 inverted microscope, equipped with a hardware autofocus system (Perfect Focus System 4) and a CrestOptics X-Light V3 spinning-disk confocal unit. Images are acquired with a high-sensitivity Kinetix 10MP sCMOS camera (Teledyne Photometrics) through a CFI PLAN APO TIRF/DIC 60×/1.49 NA oil-immersion objective (Nikon). We excite the WGA-Alexa Fluor 555 using a 561 nm solid-state laser line, collecting the emission through a standard 595/50 nm bandpass filter. NIS-Elements software (Nikon) manages image acquisition and motorized stage control, while subsequent processing of the fluorescence images is conducted in Fiji (ImageJ).

### Image-based nematic-field analysis

We extract the local cell-axis orientation from bright-field images using a structure-tensor analysis of image gradients^66^. Because image intensity typically varies most strongly across elongated cells, we rotate the principal gradient direction by *π/*2 to obtain the local cell-axis angle *θ*(**x**). To account for nematic head–tail symmetry, we represent this orientation using the doubled-angle vector **q**(**x**) = (cos 2*θ*, sin 2*θ*). We also construct binary biomass masks *M* (**x**), with *M* = 1 in segmented diatom regions and *M* = 0 elsewhere. Mask construction and biomass-occupancy thresholds are detailed in the Supplementary Information. Director rods and line-integral-convolution textures are used for visualization.

To characterize the nematic architecture of tidal-flat diatoms, we use three complementary measures that yield three characteristic length scales. First, to quantify orientational coherence, we evaluate the radially averaged connected orientational correlation, *C*_conn_(*r*), where *r* = |Δ**x**| represents the separation distance. We fit its decay to a single exponential, *C*_conn_(*r*) = *A*_0_ exp(−*r*/*ξ*), to extract the correlation length *ξ*, with *A*_0_ as a fitted amplitude. Second, we define the local nematic order at observation scale *L* as *s*_2_(*L*, **x**) = |⟨**q**⟩_*L,M*_|, where ⟨· ⟩_*L,M*_ denotes averaging over biomass pixels (*M* = 1) within a square window of size *L* × *L* centered at **x**. We calculate *S*_2_(*L*) as the mean of *s*_2_ (*L*, **x**) over complete windows with at least 50% biomass-mask occupancy, and define 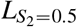 as the crossover scale at the first downward crossing of *S*_2_(*L*) = 0.5. Finally, we locate winding-validated +1/2 and −1/2 singularities in a mask-normalized, coarse-grained **q** field. We summarize their abundance using the defect-density length, *ℓ*_defect_ = (*A*_search_/*N*_defect_)^1/2^, where *A*_search_ is the area satisfying the defect-search criteria and *N*_defect_ is the total number of retained defects.

To analyze geometric distortions, we Gaussian-smooth the doubled-angle field **q** to obtain 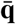. The corresponding director angle is 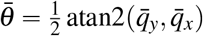, giving 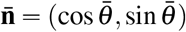. We then calculate the two-dimensional splay, 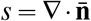, and bend, 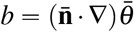^67, 68^. We compare dimensionless, local-order-weighted deformation scores derived from *s*^2^ and *b*^2^. Coarse-graining scales, defect validation and deformation-score definitions are detailed in the Supplementary Information.

### Trigger-wave, cell-motion and alignment remodeling analysis

We quantify the dynamics of the trigger waves by integrating wave-region segmentation, nematic-guided wavefront tracking, individual-cell trajectories, and particle image velocimetry (PIV). This combined approach allows us to capture the motion of individual cells (via trajectories), the propagation speed of trigger waves (via wave front tracking), and the Eulerian velocity of the local diatom population in trigger waves (via PIV).

For wave-region segmentation, we identify candidate wave-region pixels by looking for locations showing a simultaneous drop in PIV cross-correlation and increase in frame-to-frame intensity changes at each analyzed time point. After filtering out small disconnected components and image-edge artifacts, we retain the dominant connected region. By accumulating these spatial masks over time, we obtain the wave-swept region, *M*_wave_(**x**,*t*). The first analyzed time at which a position enters the wave-swept region marks the wave-arrival time. We further extract the advancing wavefront from the region’s outer boundary.

To visualize wave-associated collective motion, we construct a historical PIV velocity map by retaining, at each wave-swept position, the velocity vector recorded when the local PIV speed is maximal. After smoothing this map, we propagate virtual tracers through the corresponding unit-vector field. These tracers are used only for visualization, not for calculating wavefront speed. Separately, we measure local wavefront speed by tracing paths between successive wavefront contours along the local nematic director field. Dividing each path length by the interval between contours gives the local director-guided propagation speed. Details of wave segmentation, tracer visualization and front-speed measurement are provided in the Supplementary Information.

We extract individual-diatom trajectories using TrackMate^69^. To categorize diatom states, we assign trajectory intervals to the ‘rested’ or ‘activated’ state when both endpoints lie outside or inside the time-dependent wave-swept region *M*_wave_ at their respective times. We then use these categorized segments to calculate diatom speeds and mean-squared displacements. Speeds are averaged separately for each trajectory and state. Similarly, we average the mean-squared-displacement values at each time lag first within individual trajectories, and subsequently across the entire trajectory ensemble.

To investigate how wave-associated population motion relates to diatom density and nematic organization, we first estimate a relative biomass-density proxy from background-corrected bright-field transmission. We calculate *ρ*_*I*_ = max[0, ln(*I/B*)], where *I* is the measured bright-field intensity and *B* is a smooth illumination background. We then normalize and locally average this signal within each dataset to obtain the normalized density field *ρ*_norm_. Next, we spatially register near-front PIV measurements from the activated side of the wave to the pre-wave density and nematic fields. We calculate the median PIV speed within common density–nematic order bins for each dataset and average the resulting maps across datasets. Within each dataset, we calculate weighted Spearman correlations to quantify the associations of binned median PIV speed with density and nematic order separately, using the number of measurements in each bin as its weight. Finally, to assess directional guidance of wave-associated population motion, we calculate the axial mismatch Δ*ϕ* = arccos(|cos(*ϕ*_velocity_ − *ϕ*_director_|), where *ϕ*_velocity_ is the PIV velocity angle and *ϕ*_director_ is the local pre-wave nematic director angle.

For recurrent waves, we measure the mean nematic order and effective density-occupied area fraction within the final cumulative wave-swept region. The effective density-occupied area fraction quantifies how broadly the density signal is distributed: it approaches one for a uniform distribution across the region and decreases as the density becomes concentrated within a smaller area. Separately, to quantify recovery of the nematic pattern after trigger-wave passage, we calculate the zero-shift correlation between the current field and the initial pre-wave reference over their common valid director support. The recovery time extends from the post-wave correlation minimum to the fitted onset of the subsequent plateau.

We assess topological remodeling by matching same-charge defects one-to-one to a reference catalogue established in the first analyzed frame, using nearest-neighbor matching within 126 *µ*m. We classify associated defects as ‘matched’, unmatched defects in the current frame as ‘emerged’, and unmatched defects in the reference frame as ‘disappeared’. To map persistent director-field remodeling between two relaxed pre-wave states, we compute the axis-folded mismatch Δ*θ*(**x**) = arccos[|cos(*θ*_2_(**x**) − *θ*_1_(**x**))|], where *θ*_1_ and *θ*_2_ represent the local director angles of the two states. We provide detailed sampling rules and quality-control criteria in the Supplementary Information.

### Numerical models of wave routing and nematic memory

To investigate the mechanisms driving trigger wave dynamics in the nematic mosaics of tidal-flat diatoms, we use two complementary numerical models that address distinct physical questions. First, we develop a reduced stochastic model to test whether the experimentally measured pre-wave director and density fields can account for the observed wave routing. In this framework, branching propagation elements move across a static director field. At each algorithmic step, an element selects the branch of the local director most closely aligned with its previous heading. This orientational guidance is combined with weak directional persistence, and the element advances subject to additive positional noise. Local cell density determines the stopping probability through a logistic function, making low-density regions less permissive to wave passage. As stochastic branching broadens the propagation footprint, a visitation-time rule imposes a transient refractory state that limits repeated passage through recently swept regions.

We compare ‘alignment-only’ and ‘density-gated’ conditions using identical, experimentally measured director and density fields, initiation sites, and random seeds. For each condition, we run a stochastic realization of 1,000 steps, accumulating the positions reached by surviving elements into a cumulative visitation map. The resulting simulated footprints are then compared against the experimentally observed wave-swept regions. Model equations, parameter values and implementation details are provided in the Supplementary Information.

Second, to capture the dynamic interplay between the trigger wave, cellular alignment, and the mucilage matrix, we construct a two-dimensional phenomenological continuum model. This model tracks multiple coupled variables: cell density, FitzHugh–Nagumo-type excitable variables (activator and recovery), an activation cue, cellular nematic order, a cellular director field, and a non-advected mucilage matrix director field. In this system, activated cells produce and transport the cue. As cue levels rise, they promote cellular motility, weaken pinning to the matrix, and drive active cell flux along the local nematic axis, with polarity selected to favor motion down the cue gradient. Concurrently, the scalar cellular nematic order relaxes toward a target value supported by attachment to the local mucilage matrix. The two director fields in this continuum model evolve on distinct timescales. The cellular director evolves through advection and relaxation of spatial variations in orientation, and realigns with the matrix after activation. The matrix director evolves more slowly and is rewritten toward the cellular orientation during the activated state.

Simulations are initialized with +1/2 and −1/2 defect textures to examine defect-mediated wave propagation, and with a −1/2 texture to examine nematic-memory remodeling. The continuum equations are integrated on a 300 × 300 grid using conservative upwind discretization for advective transport, a standard five-point stencil for the Laplacian and zero-gradient boundary conditions. Recovery and remodeling are quantified by cellular and matrix orientation correlations relative to their initial states. Model equations, initial conditions and simulation protocols are described in the Supplementary Information.

### External stimulation and patterned-substrate fabrication

To test external control of trigger-wave initiation, we subject dense *Cylindrotheca* sp. mats to localized mechanical, chemical, and optical perturbations. We perform mechanical and chemical stimulation using a micromanipulation and microinjection setup (Scientifica and Sutter Instrument). We fabricate the micropipettes from filamented borosilicate glass capillaries (1B100F-4; World Precision Instruments) using a P-97 Flaming/Brown micropipette puller (Sutter Instrument). The puller is configured with a filament-heat setting of 757, a hard-pull setting of 50, a velocity trip point of 70, a 200-ms delay, and a cooling-air pressure setting of 500. We subsequently bend the micropipettes using an MF-900 microforge (Narishige). For mechanical stimulation, we lower a pulled glass micropipette (approximately 3-5 *µ*m inner tip diameter) under bright-field observation until it gently contacts and locally indents the biofilm surface.

For chemical stimulation, we dissolve *trans,trans*-2,4-decadienal (technical grade, 85%; CAS 25152-84-5; MilliporeSigma) in anhydrous methanol to prepare a 10 mM stock solution. We then dilute this stock 1:1,000 in filtered seawater to achieve a final concentration of 10 *µ*M as an injection solution. We load the solution into a glass micropipette and use a Scientifica micromanipulator to position its tip immediately above the diatom biofilm without contacting the cells. During injection, we apply 10 successive pressure pulses using a XenoWorks Digital Microinjector (Sutter Instrument) operated in pulse mode (40 hPa for 0.03 s per pulse), while maintaining a constant transfer/compensation pressure of +9 hPa.

For optical stimulation, we illuminate a user-defined circular region of interest using a Nikon Eclipse Ti2 inverted microscope equipped with a CrestOptics X-Light V3 spinning-disk unit and an iLAS2 FRAP/photostimulation module. We apply a single scan using a 473-nm laser at 20% nominal power with a 10 *µ*s pixel dwell time, and record the resulting trigger-wave response on the same microscope through a 20×/0.8 NA objective.

To prescribe the cellular alignment field, we fabricate patterned SU-8 masters via maskless photolithography. Silicon wafers (UniversityWafer) are cleaned in a Diener Nano plasma asher (Diener electronic) using oxygen plasma at a power of 300 W for 300 s. The wafers are then coated with SU-8 2010 photoresist by spin coating at 2,000 rpm for 30 s using a WS-650MZ spin coater (Laurell Technologies). After a 3-minute soft bake at 95 °C, we expose the wafers to computer-designed patterns at 365 nm using an ML3 MicroWriter direct-write lithography system (Durham Magneto Optics) with an exposure dose of 128 mJcm^−2^. Following exposure, we post-bake the wafers at 95 °C for 3 min and develop them in SU-8 developer for 4 min. We verify the resulting SU-8 relief height to be 11–12 *µ*m using an Alpha-Step stylus profilometer (KLA-Tencor), and subsequently use these patterned wafers as masters for polydimethylsiloxane (PDMS) replication.

We mix PDMS base and curing agent (SYLGARD 184 Silicone Elastomer Kit) at a 10:1 mass ratio, degas the mixture, and cast it over the SU-8 master. After curing the PDMS at 60 °C for 2 hours, we peel it from the master to produce a negative replica containing the patterned surface grooves. The replicas are mounted in a petri dish and used as the final culture substrates. Prior to *Cylindrotheca* sp. seeding, we plasma-treat the PDMS substrates and immerse them in f/2 medium for 1 hour. Diatoms are cultured for 1-2 weeks before imaging.

## Data availability

The data supporting the findings of this study are available within the text, Extended Data, Supplementary Information, and [Source Data files](https://doi.org/10.5281/zenodo.22573663).

## Code availability

The code used in this study is available on [Zenodo](https://doi.org/10.5281/zenodo.22573663).

## Acknowledgements

We thank N. Hall for assistance on the tidal-flat sampling approach. We thank J. Chen for assistance on the micro-injection protocol. We thank all members of the Prakash lab for discussions. Part of this work was performed at the Cell Sciences Imaging Facility (RRID: SCR_017787) and the Stanford Nano Shared Facilities (RRID: SCR_026695) at Stanford University, and we thank D. Lenzi and N. Reihaneh for their assistance. We thank the Electron Microscope Lab at the University of California, Berkeley and the assistance provided by M. Kang. Q.Z. and M.P. acknowledge support from Human Frontier Science Program and National Science Foundation (NSF) (grant no. 135316). M.P. acknowledges further financial support from the Schmidt Futures Innovation Fellowship, the Moore Foundation, the Dalio Foundation, the NSF Center for Cellular Construction (grant no. DBI-1548297), ARIA, and the Woods Institute for the Environment.

## Author contributions

Q.Z. and M.P. designed the research. Q.Z. and M.P. performed field studies. Q.Z. performed experiments. In discussion with M.P., Q.Z. analysed the data and performed theoretical and computational analyses. Q.Z. and M.P. wrote the manuscript.

## Additional information

**Supplementary information** is available for this paper.

