## Supplemental Information for "Diatom Logic: Rewritable active nematic circuits in tidal-flat ecosystems"

**The PDF file includes:**

Supplementary Information

Figs. S1 to S10

Table. S1

Captions for Supplementary Movies 1 to 7

**Other Supplementary Information for this manuscript includes the following:**

Supplementary Movies 1 to 7

### Local alignment recurs in native tidal-flat biofilms

We collect intact surface blocks from Aardvark Beach in San Mateo and the Duck Pond site near Palo Alto, California. Each block measures approximately $10 \times 10$ cm across and $2$ cm deep. We keep the surface biofilm intact during transfer and image it within 1–2 h of collection.

Images from February, March and September 2025 show abundant elongated pennate diatoms arranged in locally aligned patches (Fig. S1). The alignment axis varies across each field, forming a heterogeneous orientational mosaic rather than a globally parallel array. The recurrence of this organization across two sites and multiple sampling dates indicates that local alignment is not confined to a single field sample or generated only after laboratory cultivation.

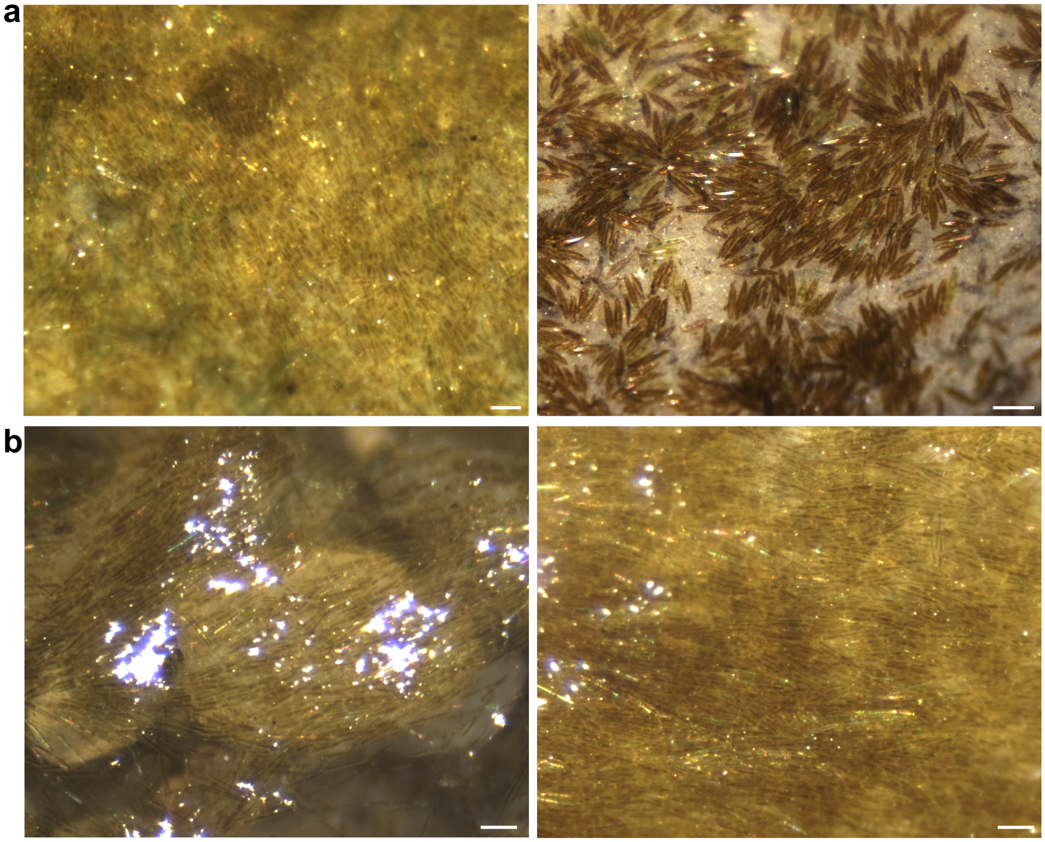


**Fig. S1. Local alignment recurs in intact tidal-flat surface biofilms across sites and sampling dates. a** Representative surface-layer micrographs from the Duck Pond site near Palo Alto, California, collected in March (left) and February (right) 2025. **b** Representative surface-layer micrographs from Aardvark Beach, San Mateo, California, collected in September 2025. Across these fields, elongated pennate diatoms form locally aligned patches whose preferred axes vary spatially across the biofilm. Scale bars, 50 $\mu$m.

### A second tidal-flat diatom isolate reconstitutes nematic organization

To test whether nematic reconstitution is restricted to the *Cylindrotheca* sp. isolate used for quantitative experiments, we examine a second tidal-flat morphotype, provisionally identified as *Navicula* sp. We use the same isolation and cultivation procedures for both isolates.

The *Navicula* sp. isolate also forms dense, orientationally ordered mats in culture (Fig. S2). Neighboring cells align into curved bundles that merge, diverge and change orientation across larger fields of view. The resulting multiscale nematic mosaic is similar to the static architecture formed by *Cylindrotheca* sp. These observations show that nematic self-organization occurs in more than one tidal-flat diatom morphotype.

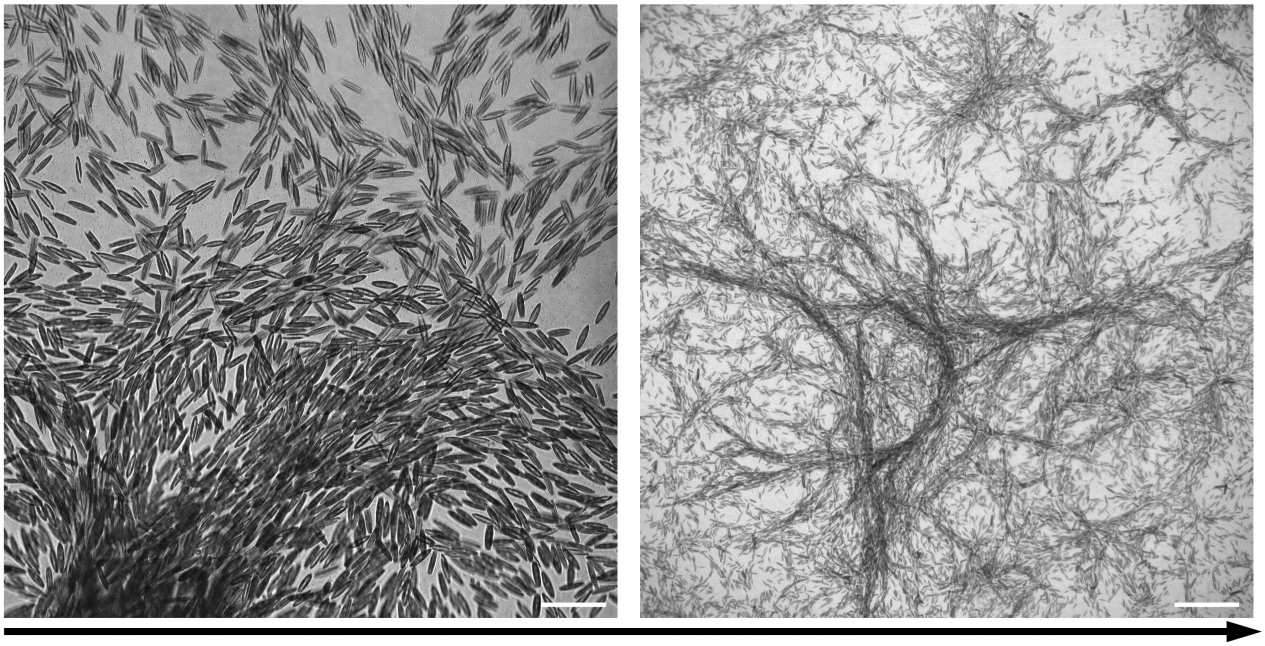

**Fig. S2. A second tidal-flat diatom isolate self-organizes into a multiscale nematic texture.** Representative bright-field micrographs of a tidal-flat morphotype provisionally identified as *Navicula* sp., shown at increasing fields of view from left to right. Neighboring cells align into curvilinear bundles; across the wider field, these bundles merge, diverge, and change orientation to form a spatially heterogeneous nematic mosaic. The arrow denotes increasing field of view. Scale bars, 50 $\mu$m (left) and 200 $\mu$m (right).

### A hierarchy of nematic length scales in a defect-rich mosaic

We characterize the nematic mosaic of tidal-flat diatoms using three complementary measures: orientational correlations, scale-dependent nematic order and defect density. These yield the characteristic lengths: the orientational correlation length $\xi$, the half-order scale $L_{S_2=0.5}$, and the defect-density length $\ell_{\text{defect}}$, respectively.

**Director-field extraction and biomass masks.** We extract local cell-axis orientations from greyscale bright-field images, with intensity $I(\mathbf{x})$ normalized to the range 0–1. From the image gradients $G_x$ and $G_y$, we construct the locally averaged structure tensor[1],

$$\mathbf{J}(\mathbf{x}) = \begin{pmatrix} J_{xx} & J_{xy} \\ J_{xy} & J_{yy} \end{pmatrix} = \begin{pmatrix} \langle G_x^2 \rangle_{\sigma_\theta} & \langle G_x G_y \rangle_{\sigma_\theta} \\ \langle G_x G_y \rangle_{\sigma_\theta} & \langle G_y^2 \rangle_{\sigma_\theta} \end{pmatrix}, \tag{S1}$$

where $\langle \cdot \rangle_{\sigma_\theta}$ denotes Gaussian smoothing with standard deviation $\sigma_\theta = 1.25\ \mu\text{m}$ for the $20\times$ and $40\times$ images, and $6\ \mu\text{m}$ for the $4\times$ images. We rotate the principal gradient direction by $\pi/2$ to obtain the local cell-axis angle: $\theta(\mathbf{x}) = \frac{1}{2}\,\text{atan2}\,(2J_{xy}, J_{xx} - J_{yy}) + \frac{\pi}{2}$. The nematic director is $\mathbf{n} = (\cos\theta, \sin\theta)$, where $\mathbf{n} \equiv -\mathbf{n}$, and its doubled-angle representation is $\mathbf{q} = (\cos 2\theta, \sin 2\theta)$.

We construct a binary biomass mask $M$ from the original greyscale image $I$, with $M = 1$ in segmented diatom regions and $M = 0$ elsewhere. We identify cell-occupied regions from intensity contrast in the $20\times$ and $4\times$ images and from local texture in the $40\times$ director overlays. For the $20\times$ images, we identify dark diatom regions using the intensity cutoff $I < 0.45$. For the $4\times$ images, locally adaptive thresholds identify dark regions relative to the surrounding mean intensity[2], and the sensitivity, which controls how readily pixels are included in the biomass mask, is fixed at $0.88$ across images. For the $40\times$ director overlays, we use local intensity variation to distinguish textured cell-occupied regions from smoother background. We quantify this variation using the local standard deviation of image intensity, then smooth and threshold the resulting map. Furthermore, to suppress segmentation artifacts, we refine the biomass masks by bridging narrow gaps, filling small background holes and removing small isolated regions.

For visualization, we represent local directors by short line segments at regularly spaced grid points within the biomass mask (Fig. S3a). We omit positions with low structure-tensor coherency and scale segment lengths within each image in proportion to $\gamma(\mathbf{x}) = \frac{\lambda_{\max} - \lambda_{\min}}{\lambda_{\max} + \lambda_{\min}}$, where $\lambda_{\max}$ and $\lambda_{\min}$ are the eigenvalues of $\mathbf{J}$. Line integral convolution (LIC) provides a continuous visualization of the coarse-grained director field (Fig. S3b).

**Scale-dependent nematic order.** To determine how far local alignment persists across the mosaic, we measure nematic order in square observation windows of increasing size. For a window of size $L \times L$ centered at $\mathbf{x}$, the local order is

$$s_2(L, \mathbf{x}) = \left|\langle \mathbf{q} \rangle_{L,M}\right| = \sqrt{\langle q_x \rangle_{L,M}^2 + \langle q_y \rangle_{L,M}^2}, \tag{S2}$$

where $\langle \cdot \rangle_{L,M}$ denotes averaging over biomass pixels within the window. We obtain the mean scale-dependent order, $S_2(L)$, by averaging $s_2(L, \mathbf{x})$ over windows that lie within the image and contain biomass over at least $50\%$ of their area. We define the half-order scale $L_{S_2=0.5}$ as the window size at which $S_2(L)$ first falls to $0.5$ as $L$ increases, using linear interpolation in $\log L$.

**Orientational correlation length.** To quantify the distance over which director orientations remain correlated across the nematic mosaic, we calculate the spatial autocorrelation of the doubled-angle field within the biomass mask. For a displacement vector $\Delta\mathbf{x}$, the raw correlation is defined as

$$C_{\text{raw}}(\Delta\mathbf{x}) = \frac{\sum_{\mathbf{x}} M(\mathbf{x})M(\mathbf{x}+\Delta\mathbf{x})\mathbf{q}(\mathbf{x}) \cdot \mathbf{q}(\mathbf{x}+\Delta\mathbf{x})}{\sum_{\mathbf{x}} M(\mathbf{x})M(\mathbf{x}+\Delta\mathbf{x})}, \tag{S3}$$

where $M$ is the biomass mask. This normalization accounts for the changing number of valid biomass-pixel pairs at different displacements. We obtain the radial correlation $C_{\text{raw}}(r)$ by grouping biomass-pixel pairs into narrow intervals of separation distance $r = |\Delta\mathbf{x}|$ and averaging their orientational correlations. Displacements with too few contributing biomass-pixel pairs are excluded.

When global nematic order is nonzero, the raw correlation can approach $S^2$ rather than zero at large separations, where $S = |\langle \mathbf{q} \rangle_M|$ is the global nematic order within the biomass mask. To isolate spatial correlations beyond this global alignment, we define the connected correlation as

$$C_{\text{conn}}(r) = \frac{C_{\text{raw}}(r) - S^2}{1 - S^2}. \tag{S4}$$

Subtracting $S^2$ removes this baseline, while division by $1 - S^2$ normalizes the correlation to one at zero separation.

We fit the decay of the connected correlation with a single exponential, $C_{\text{conn}}(r) = A_0 \exp\left(-\frac{r}{\xi}\right)$, where $A_0$ is the fitted amplitude, and $\xi$ is the orientational correlation length. Table S1 reports the fitted amplitude $A_0$, correlation length $\xi$, $R^2$, realized fit range, and number of fitted bins for each independent dataset. Fits whose maximum analyzed separation is less than $2\xi$ are marked as field-of-view (FOV) limited. The stitched $20\times$ field and all $4\times$ fields sample more than two correlation lengths, whereas the three individual $20\times$ fields indicate this limitation. We retain the results from $20\times$ for comparison, considering that they are consistent with the correlation lengths obtained from larger-FOV $4\times$ and stitched $20\times$ measurements.

**Defect-density length.** We identify $+1/2$ and $-1/2$ defects in the coarse-grained nematic field and convert their number per unit area into a characteristic length. We first average the doubled-angle field over biomass using a radially symmetric triangular kernel[3],

$$\overline{\mathbf{q}}(\mathbf{x}) = \frac{[K_R * (M\mathbf{q})](\mathbf{x})}{[K_R * M](\mathbf{x})}, \qquad K_R(d) \propto \begin{cases} R - d, & 0 \le d < R, \\ 0, & d \ge R. \end{cases} \tag{S5}$$

Here, $*$ denotes spatial convolution, $d$ is the distance from the kernel center and $R = 80\ \mu\text{m}$ is the averaging radius. The value is selected to suppress fine-scale variations while preserving the larger-scale organization of the nematic mosaic. The kernel weights decrease linearly with distance and sum to one. The denominator is therefore the weighted fraction of the neighborhood occupied by biomass; we retain positions where this fraction exceeds $20\%$ to avoid poorly supported orientation estimates.

Candidate defect cores are local minima of $|\overline{\mathbf{q}}|$, located after Gaussian smoothing with standard deviation $8\ \mu\text{m}$ to suppress small spurious minima. We retain candidate cores satisfying $|\overline{\mathbf{q}}| \le 0.75$. To evaluate the charge locally, we follow the doubled-angle phase $\varphi = \text{atan2}(\bar{q}_y, \bar{q}_x)$ around a square contour centered on each candidate core, with each side approximately $25\ \mu\text{m}$ from the core. The nematic charge follows from the phase winding[4],

$$s = \frac{\Delta_{\mathcal{C}}\varphi}{4\pi}, \tag{S6}$$

where $\Delta_{\mathcal{C}}\varphi$ is the total unwrapped phase change accumulated in one traversal of the contour. We accept a candidate as a $+1/2$ or $-1/2$ defect if its calculated charge lies within $0.1$ of the corresponding value.

A broad defect core can produce several candidate minima. To avoid repeated counting, we group same-sign candidates by linking pairs separated by at most $62.5\ \mu\text{m}$. Within each of these groups, we retain the candidate with the smallest $|\overline{\mathbf{q}}|$. Opposite-sign candidates remain separate. The defect markers shown in Fig. S3b correspond to the winding-validated defects retained by this procedure.

To reduce boundary effects, we exclude positions within $60\ \mu\text{m}$ of the image edge. The searchable area, $A_{\text{search}}$, comprises the remaining positions where kernel-weighted biomass occupancy exceeds $20\%$ and the surrounding square winding contour lies entirely within the region satisfying this threshold. We calculate the defect-density length as

$$\ell_{\text{defect}} = \rho_{\text{defect}}^{-1/2} = \sqrt{\frac{A_{\text{search}}}{N_{\text{defect}}}}, \tag{S7}$$

where $N_{\text{defect}}$ is the total number of retained $+1/2$ and $-1/2$ defects within this area.

We measure $\xi$ and $L_{S_2=0.5}$ in three individual $20\times$ fields, one stitched $20\times$ field and four $4\times$ fields ($n = 8$). Defect-density measurements use the stitched $20\times$ field and the four $4\times$ fields ($n = 5$). Each field contributes one estimate, and we report the mean $\pm$ standard error of the mean (s.e.m.) across fields.

### Positional correlations in tidal-flat diatom nematic mats

To test whether locally aligned diatoms form periodic spatial arrangements, we calculate director-aligned pair-distribution functions separately for four independent datasets: one $10\times$ field, two stitched $20\times$ fields and one stitched $40\times$ field. Within each analyzed image patch of approximately $120 \times 120\ \mu\mathrm{m}^2$, we project cell-center displacements along and across the patch-averaged nematic director:

$$r_{\parallel,ij} = (\mathbf{r}_j - \mathbf{r}_i) \cdot \mathbf{n}, \qquad r_{\perp,ij} = (\mathbf{r}_j - \mathbf{r}_i) \cdot \mathbf{n}_\perp, \tag{S8}$$

where $\mathbf{r}_i$ is the center of cell $i$, $\mathbf{n}$ is the patch-averaged director and $\mathbf{n}_\perp$ is the perpendicular unit vector. We exclude self-pairs and count each pair in both displacement directions.

We generate a random reference by placing the same number of cell centers uniformly within each full rectangular patch and projecting their displacements onto the same director axes. Pair displacements are counted in two-dimensional bins of width 2 $\mu$m along each axis, with bin centers spanning $-60$ to 60 $\mu$m. For each dataset, we sum the observed and randomized counts separately across patches before calculating the pair-distribution function:

$$g(r_\parallel, r_\perp) = \frac{H_{\mathrm{data}}(r_\parallel, r_\perp)}{\langle H_{\mathrm{random}}(r_\parallel, r_\perp) \rangle}. \tag{S9}$$

Here, $H_{\mathrm{data}}$ and $H_{\mathrm{random}}$ are the observed and randomized pair counts in each bin, respectively, and $\langle \cdot \rangle$ denotes averaging over repeated randomizations. Thus, $g = 1$ indicates the pair frequency expected from random placement.

The stitched $40\times$ map shows an elongated depletion of pairs at short separations, consistent with anisotropic short-range packing (Fig. S3c). For each dataset, we obtain the parallel and perpendicular profiles by summing the observed and reference pair counts separately within 5 $\mu$m of the corresponding axis and taking their ratio. Across all four datasets, these profiles approach $g = 1$ without sustained, regularly spaced peaks (Fig. S3d). Together with the orientational measurements above, these results support local nematic alignment without detectable periodic positional order over the measured range.

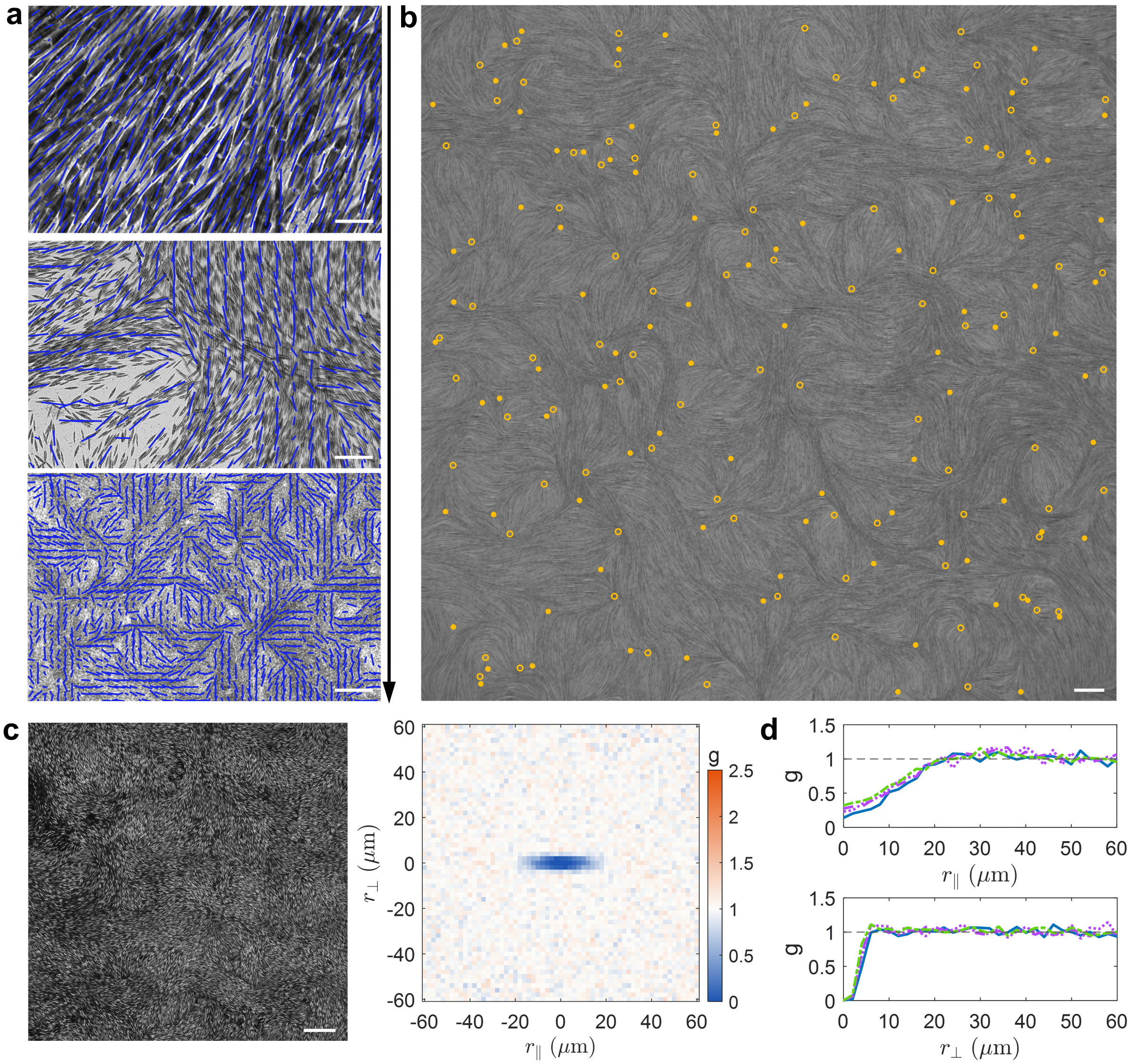


**Fig. S3. Positional correlations and multiscale nematic organization.** (**a**) Representative bright-field images overlaid with nematic directors (blue) extracted by structure-tensor analysis. The overlays reveal cell-scale alignment, curved bundles and orientational domains over increasing fields of view (top to bottom; arrow). Scale bars, 50, 100 and 500 $\mu$m, respectively. (**b**) Coarse-grained director texture rendered by line-integral convolution (LIC), revealing the domain-scale organization and associated $\pm 1/2$ topological defects. Open and filled yellow circles denote $+1/2$ and $-1/2$ defects, respectively. Scale bar, 200 $\mu$m. (**c**) Representative stitched $40\times$ image (left) and corresponding director-aligned cell-center pair-distribution function $g(r_\parallel, r_\perp)$ (right). Color denotes $g$. Scale bar, 200 $\mu$m. (**d**) Pair-correlation profiles parallel (top) and perpendicular (bottom) to the patch-averaged director from four datasets: one $10\times$ field (blue), two stitched $20\times$ fields (purple) and one stitched $40\times$ field (green). Gray dashed lines mark the random-reference level, $g = 1$.

### The curvilinear nematic mosaic is splay-rich at the mesoscale

Splay describes the local convergence or divergence of director lines, whereas bend describes their curvature. We compare their spatial distributions and mean deformation scores to quantify the relative contributions of these modes to the mesoscale nematic mosaic of tidal-flat diatoms.

We smooth the doubled-angle field $\mathbf{q}$ with a Gaussian kernel of standard deviation $\ell_d = 60\ \mu\mathrm{m}$: $\overline{\mathbf{q}} = G_{\ell_d} * \mathbf{q}$, and define $Q = |\overline{\mathbf{q}}|$. The magnitude $Q$ measures local orientational order. The corresponding director angle and director are $\overline{\theta} = \frac{1}{2}\,\mathrm{atan2}\left(\overline{q}_y, \overline{q}_x\right)$ and $\overline{\mathbf{n}} = (\cos\overline{\theta}, \sin\overline{\theta})$. For these densely populated images, we analyse the continuous director field without a biomass mask to avoid artificial gradients at mask boundaries.

We estimate angular derivatives from the smoothed doubled-angle field:

$$\partial_i\overline{\theta} = \frac{1}{2}\frac{\overline{q}_x\,\partial_i\overline{q}_y - \overline{q}_y\,\partial_i\overline{q}_x}{\max\left(Q^2, Q^2_{\mathrm{min}}\right)}, \qquad i = x, y. \tag{S10}$$

The lower bound $Q_{\mathrm{min}} = 0.15$ limits numerical amplification where the local orientation is poorly defined.

Using these angular derivatives, we evaluate the two-dimensional splay and bend fields based on their standard geometric definitions[5]:

$$\begin{aligned} s &= -\sin\overline{\theta}\,\partial_x\overline{\theta} + \cos\overline{\theta}\,\partial_y\overline{\theta}, \\ b &= \cos\overline{\theta}\,\partial_x\overline{\theta} + \sin\overline{\theta}\,\partial_y\overline{\theta}. \end{aligned} \tag{S11}$$

These estimate the geometric deformations $s = \nabla\cdot\overline{\mathbf{n}}$ and $b = (\overline{\mathbf{n}}\cdot\nabla)\overline{\theta}$. We compare them using dimensionless, local-order-weighted scores: $D^{(0)}_{\mathrm{splay}} = Q^2(\ell_d s)^2$ and $D^{(0)}_{\mathrm{bend}} = Q^2(\ell_d b)^2$. The factor $Q^2$ reduces contributions from weakly ordered regions, while $\ell_d$ expresses both deformations relative to the same spatial scale. We then smooth the score maps with a narrower Gaussian kernel of standard deviation $\sigma_D = 20\ \mu\mathrm{m}$ to reduce residual small-scale variation: $D_{\mathrm{splay}} = G_{\sigma_D} * D^{(0)}_{\mathrm{splay}}$, and $D_{\mathrm{bend}} = G_{\sigma_D} * D^{(0)}_{\mathrm{bend}}$. These final scores are used for both visualization and quantitative analysis.

For visualization, the combined maps show splay in red, bend in cyan and their overlap in white (Figs. 1i and S4a). Both components share a display maximum set to the 97th percentile of their combined score values within each image. In the separate component maps, this maximum is calculated for each component independently to reveal its spatial pattern (Fig. S4b). These display settings do not affect the quantitative analysis. The maps reveal extended splay-rich regions where neighboring director bundles converge or diverge, including along some interfaces between orientational domains. Bend-rich regions follow strongly curved director paths. Each of the seven independent biological experiments contributes one image-level mean to the comparison of splay and bend scores (Fig. 1k).

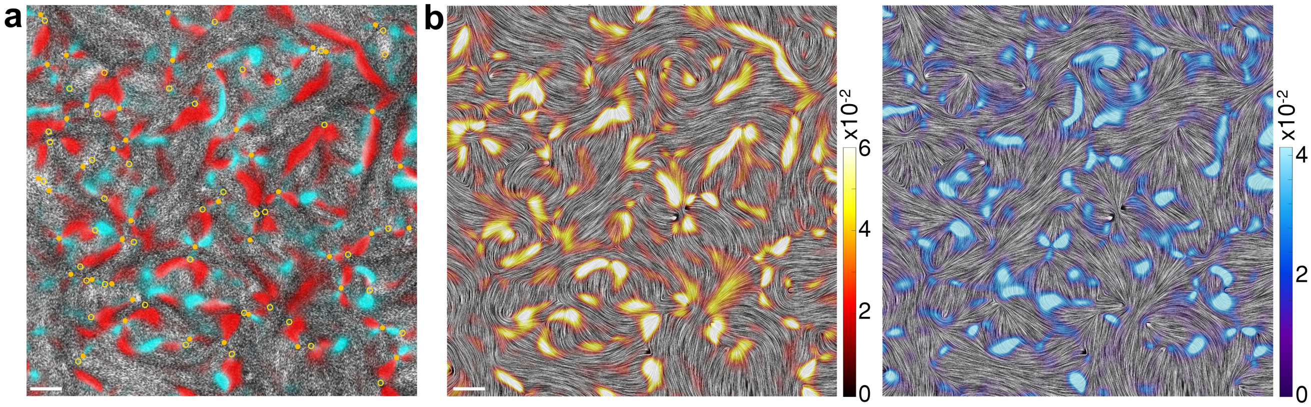


**Fig. S4. Spatial distributions of splay and bend in the nematic mosaic.** (**a**) Composite map of the dimensionless splay score $D_{\mathrm{splay}}$ (red) and bend score $D_{\mathrm{bend}}$ (cyan) over the bright-field image; overlap appears white. Open and filled yellow circles denote $+1/2$ and $-1/2$ defects, respectively. (**b**) Splay (left, warm colors) and bend (right, cool colors) shown separately over line-integral-convolution renderings of the same coarse-grained director field. Color bars show the corresponding dimensionless scores, with separate scales for splay and bend. Scale bars, 300 $\mu$m.

### Trigger-wave propagation and sequential cellular recruitment

Trigger-wave passage mobilizes a largely arrested diatom layer. To distinguish sequential recruitment from transport by a fixed group of moving cells, we compare wavefront advance with population flow and individual-cell motion. Particle image velocimetry (PIV) estimates local population velocity at fixed image positions, TrackMate follows individual cells, and wavefront tracking measures the propagation speed of the activation boundary.

**Wave-region and wavefront segmentation.** PIVlab[6, 7] provides the local population velocity $\mathbf{V}_{\mathrm{p}}(\mathbf{x}, t) = (u_{\mathrm{p}}, v_{\mathrm{p}})$ and the PIV cross-correlation coefficient $C_v(\mathbf{x}, t)$. Wave-associated rearrangement disrupts the image texture and reduces $C_v$, while cell redistribution produces a large frame-to-frame intensity change. We identify candidate wave regions where both changes occur. After removing small isolated regions and image-edge artifacts, we retain the dominant connected region at each analyzed time.

Combining these masks over time gives the cumulative wave-swept mask $M_{\mathrm{wave}}(\mathbf{x}, t)$, containing all positions reached by the wave up to time $t$. The first analyzed time at which a position enters this mask defines its wave-arrival time. We extract the wavefront $F(t)$ from the outer boundary of the cumulative mask, excluding image-edge artifacts and short disconnected fragments (Fig. S5a).

**Visualization of wave-associated collective motion.** To visualize collective motion during wave passage, we interpolate the PIV field onto the image grid. At each position within the final wave-swept region, we select the velocity vector recorded when the local PIV speed is highest and combine these vectors into a single map. After smoothing this map, we trace paths along the local velocity directions in fixed spatial steps. Paths begin near the initiation region and at front positions where the wave reaches new areas. These paths visualize wave-associated collective motion rather than trajectories of individual cells (Fig. 2a).

**Wavefront propagation speed.** We measure front advance along the local nematic direction between successive analyzed times. The region added to the cumulative mask between $t$ and $t + \Delta t$ identifies where the wave advances. From approximately evenly spaced points along $F(t)$, we trace paths along both directions of the nematic axis, $\pm\mathbf{n}$, toward the nearby portion of $F(t + \Delta t)$ associated with this advance.

Paths must connect the two local fronts and cross into the newly swept region. By contrast, we exclude paths that point inward, wind excessively or reach distant front branches, as well as paths that are too long relative to the local separation between fronts. Negligible displacements and measurements near image or analysis boundaries are also excluded. When both nematic directions meet these criteria, we select the path best aligned with the outward front normal and local direction of advance.

For each retained path, the director-guided propagation speed is

$$v_{\mathrm{wave}} = \frac{p\,\ell_{\mathrm{path}}}{\Delta t}, \tag{S12}$$

where $\ell_{\mathrm{path}}$ is the path length in pixels, $p$ is the pixel size in micrometres per pixel and $\Delta t$ is the interval between fronts. This measures propagation along the local nematic pathway. Fig. S5b shows the local speeds, and Fig. S5c shows their mean at each analyzed time. The wave-speed distribution and mean in Fig. S5e use the speeds from all retained paths across the analyzed time intervals.

**Individual-cell trajectories and gliding speed.** We track individual diatoms with TrackMate[8, 9] and match their detections to the analyzed movie frames. The speed between consecutive detections is

$$v_{\mathrm{diatoms}}(t) = \frac{|\mathbf{r}(t+\Delta t) - \mathbf{r}(t)|}{\Delta t}, \tag{S13}$$

where $\mathbf{r}(t)$ is the cell position at detection time $t$, and $\Delta t$ is the time interval. Fig. S5d shows individual-cell trajectories colored by time.

**State-resolved cell motility.** At each detection time, we classify a tracked position as rested when it lies outside the cumulative wave mask $M_{\mathrm{wave}}(\mathbf{x}, t)$ and activated when it lies inside (Fig. S5a). For speed and mean-squared displacement (MSD), we include only intervals whose two endpoints belong to the same state at their respective times.

For each state, we first average cell speeds within individual trajectories and then average across trajectories (Fig. S5e). We also calculate the MSD separately for rested and activated motion:

$$\mathrm{MSD}(\tau) = \left\langle |\mathbf{r}(t+\tau) - \mathbf{r}(t)|^2 \right\rangle, \tag{S14}$$

where $\tau$ is the time lag. At each lag, we first average squared displacements within individual trajectories and then average across trajectories. We estimate effective scaling exponents from log–log fits of $\mathrm{MSD}(\tau) \sim \tau^{\alpha}$. For rested motion, we fit the first two lag points (1.40–2.81 s) and the remaining eight points (4.21–14.04 s) separately. Activated motion is fitted over all ten lag points (1.40–14.04 s), as shown in Fig. 2d.

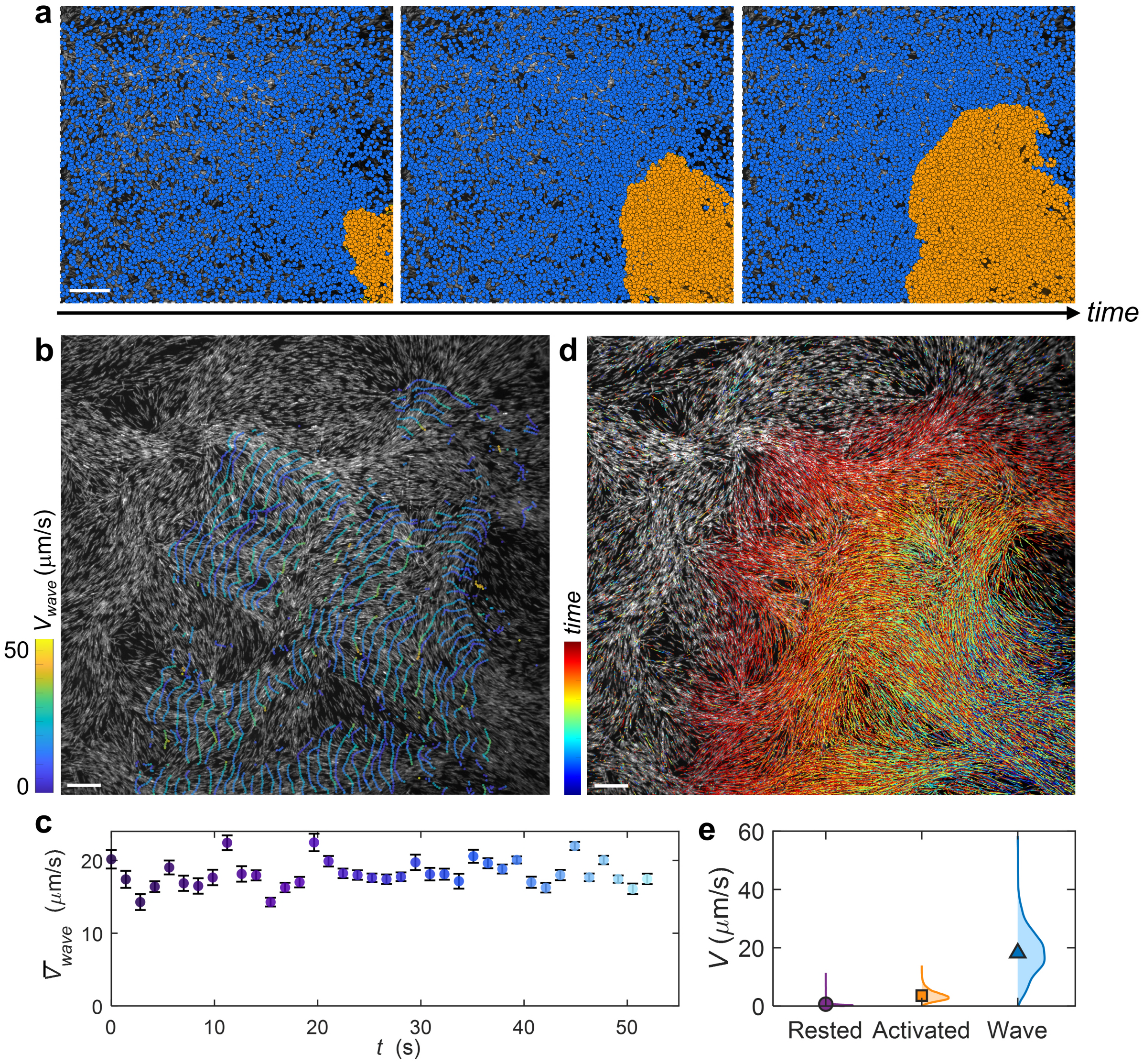


**Fig. S5. Trigger waves propagate faster than individual-cell gliding.** (**a**) Cell-state assignment using the cumulative wave-swept region: rested cells outside the region (blue) and activated cells within it (orange). (**b**) Successive segmented wavefront contours colored by the local director-guided propagation speed, $v_{\rm wave}$. (**c**) Mean wavefront speed over time; error bars show s.e.m. across local front measurements at each time. (**d**) Trajectories of individual diatoms during a representative trigger wave; color denotes time. (**e**) Half-violin distributions of per-trajectory mean speeds for rested and activated cells, compared with local wavefront speeds. Symbols denote mean. Scale bars, 200 $\mu$m (**a**) and 100 $\mu$m (**b,d**).

### Local front-speed analysis in a nonlinear reaction–diffusion–advection model

To interpret how the activation front can outrun individual activated cells, we consider a locally one-dimensional, dimensionless activation field $c(x,t)$,

$$\partial_t c + v_{\mathrm{adv}} \partial_x c = D_{\mathrm{eff}} \partial_x^2 c + r_{\mathrm{eff}} f(c), \tag{S15}$$

where $v_{\mathrm{adv}}$ is the effective directed-transport velocity, $D_{\mathrm{eff}}$ is the effective diffusivity of activation through the mat, $r_{\mathrm{eff}}^{-1}$ is the characteristic local recruitment time, and $f(c)$ represents nonlinear threshold kinetics[10–12].

For this local analysis, we assume $v_{\mathrm{adv}}$, $D_{\mathrm{eff}}$ and $r_{\mathrm{eff}}$ are constant during sustained propagation. Considering a travelling front: $c(x,t) = C(z)$, where $z = x - v_{\mathrm{front}} t$ measures position relative to the moving front along the propagation direction, Eq. (S15) becomes

$$D_{\mathrm{eff}} C'' + (v_{\mathrm{front}} - v_{\mathrm{adv}})\, C' + r_{\mathrm{eff}} f(C) = 0. \tag{S16}$$

We express $z$ in units of the length $\sqrt{D_{\mathrm{eff}}/r_{\mathrm{eff}}}$, using the dimensionless coordinate $\zeta = z\sqrt{r_{\mathrm{eff}}/D_{\mathrm{eff}}}$. The corresponding dimensionless front speed, $\alpha_{\mathrm{kin}}$, gives

$$v_{\mathrm{front}} = v_{\mathrm{adv}} + \alpha_{\mathrm{kin}} \sqrt{D_{\mathrm{eff}} r_{\mathrm{eff}}}, \tag{S17}$$

where $\alpha_{\mathrm{kin}}$ depends on the nonlinear kinetics represented by $f(c)$. We consider fronts that advance relative to the directed transport, for which $\alpha_{\mathrm{kin}} > 0$. The length $\sqrt{D_{\mathrm{eff}}/r_{\mathrm{eff}}}$ characterizes activation spreading during one recruitment time, $r_{\mathrm{eff}}^{-1}$; their ratio gives the speed scale $\sqrt{D_{\mathrm{eff}} r_{\mathrm{eff}}}$.

Under the phenomenological approximation $v_{\mathrm{adv}} \simeq v_{\mathrm{cell}}$, activated-cell motion provides directed transport, whereas the measured excess $v_{\mathrm{front}} - v_{\mathrm{cell}}$ reflects activation spreading ahead of the moving population and sequentially recruiting resting cells.

### Nematic alignment and density define accessible wave routes

To examine how the initial architecture relates to wave-associated collective motion, we compare near-front PIV velocities with density and nematic maps from the first analyzed frame of each time-lapse image sequence. These maps serve as fixed references throughout the comparison, with $\theta_0(\mathbf{x})$ denoting the local director angle in this reference frame.

**Relative biomass-density field.** Bright-field illumination varies across the image, so raw intensity reflects both biomass and background brightness. We therefore correct for uneven illumination before estimating relative biomass density. To estimate the background in each frame, we suppress dark cellular features using morphological closing with a disk of radius 252 $\mu$m. We then smooth the result with a Gaussian filter of standard deviation 94.5 $\mu$m to obtain the illumination background $B(\mathbf{x}, t)$. Dividing the original greyscale intensity $I(\mathbf{x}, t)$ by this background gives the corrected transmission: $T(\mathbf{x}, t) = \frac{I(\mathbf{x},t)}{B(\mathbf{x},t)}$. From this transmission, we define an optical-density-like signal, $\rho_I(\mathbf{x}, t) = \max[0, -\ln T(\mathbf{x}, t)]$. Regions darker relative to their local background give larger values of $\rho_I$.

For each dataset, we calculate the 1st and 99th percentiles of $\rho_I$ across sampled frames and map this range linearly to $[0, 1]$. Values below and above these limits are set to 0 and 1, respectively. We divide each image into a $500 \times 500$ grid of windows and average the normalized signal within each window to obtain the relative biomass-density score, $\rho_{\mathrm{norm}}(\mathbf{x}, t)$.

**Wave-associated population motion relative to the pre-wave architecture.** At each analyzed time, we sample PIV velocities within $\approx 160$ $\mu$m of the segmented wavefront, keeping only positions inside the cumulative wave-swept region. This selects population motion on the activated side of the front. We interpolate the reference density map to these sampling positions. At each position, we pair the PIV speed, $V_{\mathrm{p}} = \sqrt{u_{\mathrm{p}}^2 + v_{\mathrm{p}}^2}$, with the relative density $\rho_{\mathrm{norm}}$, local nematic order $S$ and director angle $\theta_0$ from the reference maps. Large cell-free regions are excluded from PIV sampling. The PIV and reference fields are shown in Figs. 3a,b.

We calculate local nematic order from the reference field: $S(\mathbf{x}) = \sqrt{\langle \cos 2\theta_0 \rangle_{\sigma_S}^2 + \langle \sin 2\theta_0 \rangle_{\sigma_S}^2}$, where $\langle \cdot \rangle_{\sigma_S}$ denotes Gaussian-weighted spatial averaging of the doubled-angle components around $\mathbf{x}$, with standard deviation $\sigma_S \approx 50$ $\mu$m. Only positions with a defined director angle contribute to the average.

For each dataset, we group measurements into bins of relative density and nematic order and calculate the median PIV speed in each bin containing data. All four independent datasets use the same bin boundaries. The resulting maps are averaged across datasets. We also calculate weighted Spearman correlations between the binned median speed and either density or nematic order, separately within each dataset. Each bin is weighted by its number of measurements, and the correlation coefficients are reported as mean $\pm$ s.e.m. across datasets (Fig. 3c).

To quantify alignment between the direction of wave-associated population motion and the initial nematic axis, we compare the PIV velocity angle, $\phi_{\mathrm{velocity}} = \mathrm{atan2}(v_{\mathrm{p}}, u_{\mathrm{p}})$, with the reference director angle at the same position, $\phi_{\mathrm{director}} = \theta_0(\mathbf{x})$. Their axial angular mismatch is

$$\Delta\phi = \arccos\left|\cos\left(\phi_{\mathrm{velocity}} - \phi_{\mathrm{director}}\right)\right| . \tag{S18}$$

Here, $\Delta\phi = 0^\circ$ denotes motion parallel or antiparallel to the director, whereas $\Delta\phi = 90^\circ$ denotes transverse motion. We normalize the joint distribution of the two angles and the distribution of $\Delta\phi$ within each dataset, then calculate the mean and s.e.m. for each distribution bin across the four datasets (Fig. 3d).

### A reduced stochastic model separates nematic guidance from density gating

Because nematic organization and biomass density vary together in the diatom mat, we use a reduced stochastic model to examine their distinct contributions to wave routing. The measured fields remain fixed: the director guides propagation direction, while density controls the probability of continued propagation.

Propagation is represented by branching elements, each describing a local propagation event rather than an individual cell. Element $i$ has position $\mathbf{x}_i^t$ and unit direction $\mathbf{e}_i^t$ at simulation step $t$. We choose the direction $\widetilde{\mathbf{n}}_i^t$ from $\pm\mathbf{n}(\mathbf{x}_i^t)$ that is closest to the element's previous direction, then update its direction as

$$\mathbf{e}_i^{t+1} = \frac{\alpha\mathbf{e}_i^t + (1-\alpha)\widetilde{\mathbf{n}}_i^t}{\|\alpha\mathbf{e}_i^t + (1-\alpha)\widetilde{\mathbf{n}}_i^t\|}, \tag{S19}$$

where $\alpha$ sets directional persistence. Its position advances according to

$$\mathbf{x}_i^{t+1} = \mathbf{x}_i^t + \Delta s\, \mathbf{e}_i^{t+1} + \sigma_D \boldsymbol{\eta}_i^t, \tag{S20}$$

where $\Delta s$ is the propagation step, $\sigma_D$ is the positional-noise amplitude and $\boldsymbol{\eta}_i^t$ is a two-dimensional standard Gaussian vector. Elements leaving the region covered by the experimental director field are removed.

We interpolate the relative-density map from the first analyzed frame onto the director-field grid, smooth it with a Gaussian filter of standard deviation $140$ px and rescale its minimum and maximum to $0$ and $1$. The resulting field, $\rho(\mathbf{x})$, determines the stopping probability at an element's new position:

$$P_{\text{stop}}(\mathbf{x}) = \frac{\lambda_\rho}{1 + \exp[k_\rho\,(\rho(\mathbf{x}) - \rho_c)]}. \tag{S21}$$

Here, $\lambda_\rho$ sets the strength of density gating, $\rho_c$ is the density at which $P_{\text{stop}} = \lambda_\rho/2$, and $k_\rho$ controls the steepness of the response. Density does not enter the direction update, and setting $\lambda_\rho = 0$ removes density gating.

Branching allows propagation to spread forward and laterally. Each surviving element branches with probability $p_b$ per step, generating five candidate daughters. Each daughter's position is given by

$$\mathbf{x}_d = \mathbf{x}_p + \ell \mathbf{e}_p + \delta_\perp \mathbf{e}_{\perp,p}, \tag{S22}$$

where $\mathbf{x}_p$ and $\mathbf{e}_p$ are the parent's position and direction, $\mathbf{e}_{\perp,p}$ is the perpendicular unit vector, and $\ell$ and $\delta_\perp$ are forward and lateral displacements. Daughters inherit the parent direction. Those landing within the director field at previously unvisited positions are accepted with probability $1 - P_{\text{stop}}(\mathbf{x}_d)$.

To represent temporary loss of excitability behind the wave, we record the most recent visit to each position. Moving elements may revisit a position within a 10-step grace interval, allowing elements within the same advancing front to overlap. They are then rejected until 60 steps have elapsed since that position's most recent visit, after which revisits are allowed again.

Each simulation starts with $4.0 \times 10^4$ elements distributed uniformly within a disk of radius 80 px near the experimental initiation site, with directions pointing radially outward. The active population is capped at $5.0 \times 10^5$ elements. Before each run, we reset the Mersenne–Twister random-number generator to seed 7. We use $\Delta s = 0.5$ px, $\sigma_D = 0.25$ px, $\alpha = 0.05$ and $p_b = 0.02$. Forward displacements are drawn uniformly from $\ell \in [5, 100]$ px and lateral displacements from $\delta_\perp \in [-5, 5]$ px. The density response uses $\rho_c = 0.65$ and $k_\rho = 10$.

We compare $\lambda_\rho = 0$ (Fig. 3e, left), $0.14$ and $0.20$ (Fig. 3e, right) with the same measured fields, initiation site and all other parameters. Each condition uses one stochastic realization of $1{,}000$ algorithmic steps. Positions reached by surviving elements form a cumulative visitation map. For visualization, each snapshot is normalized independently and shown in blue over a line-integral-convolution rendering of the static director field (Fig. S6). We compare the spatial extent and geometry of the simulated footprints across panels.

Without density gating, propagation spreads broadly along director-guided routes (Fig. S6a). Increasing $\lambda_\rho$ to $0.14$ and $0.20$ suppresses propagation through low-density regions and confines the footprint to density-supported corridors (Fig. S6b,c). The reduced model illustrates how nematic guidance and density gating shape the propagation footprint, but does not reproduce all local routing details.

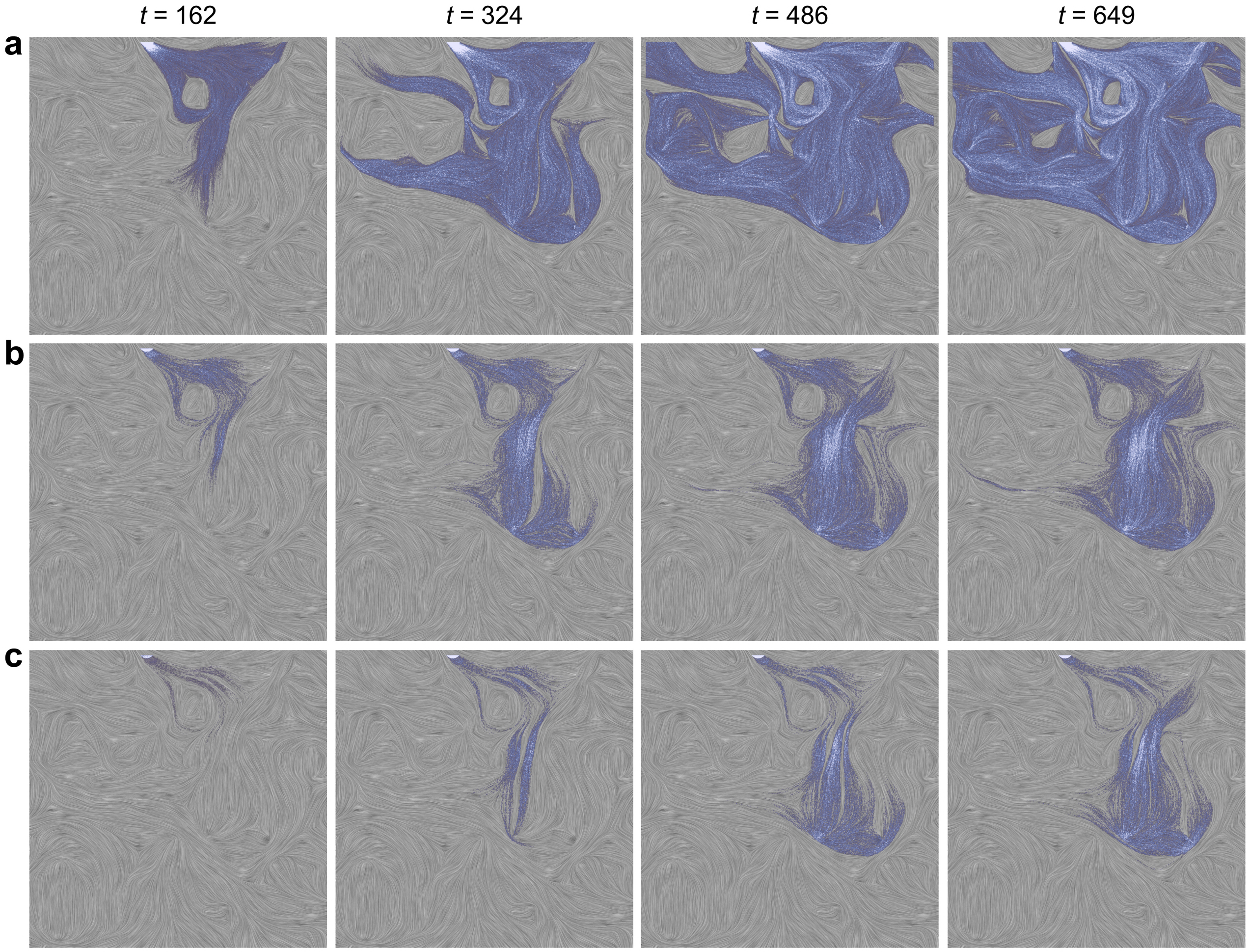


**Fig. S6. Cell density gates access to nematic-guided trigger-wave pathways.** Snapshots at increasing simulation steps $t$ (left to right), using the same measured static director and density fields. Gray line-integral-convolution textures show the director field, and blue shading shows the cumulative simulated propagation footprint, normalized independently in each snapshot. (**a**) With density gating removed ($\lambda_\rho = 0$), the footprint follows the nematic texture but spreads across a broad region. (**b,c**) Increasing density gating from $\lambda_\rho = 0.14$ (**b**) to $\lambda_\rho = 0.20$ (**c**) progressively suppresses propagation through low-density regions and confines the footprint to density-supported corridors. Each row shows one stochastic realization.

### Repeated-wave recovery and director-field evolution

To characterize the variation of nematic architecture in recurrent trigger waves, we measure both changes in mean nematic order and biomass distribution and the recovery of the spatial director pattern after repeated waves.

**Mean nematic order and biomass distribution.** We use a fixed wave-affected region, defined as the final cumulative wave-swept mask. Within this region, we calculate the area-weighted mean nematic order, $\overline{S}(t)$, and the effective density-occupied area fraction: $f_{\rho,\mathrm{eff}}(t) = \frac{\left[\sum_i w_i \rho_i(t)\right]^2}{\left(\sum_i w_i\right)\left[\sum_i w_i \rho_i^2(t)\right]}$. Here, $\rho_i(t)$ is the non-negative optical-density-like biomass signal in spatial bin $i$, and $w_i$ is the area of the fixed region contained in that bin. Values near one indicate a nearly uniform density signal across the region, whereas smaller values indicate a more uneven distribution. For visualization, we independently rescale the $\overline{S}(t)$ and $f_{\rho,\mathrm{eff}}(t)$ traces to $[0, 1]$ over the analyzed interval, and denote the rescaled traces by $S_n(t)$ and $f_{\rho,n}(t)$, respectively (Fig. 4b).

**Recovery of the spatial director pattern.** We compare each director field with the initial pre-wave field at time $t_0$. We coarse-grain the doubled-angle field, $\mathbf{q} = (\cos 2\theta, \sin 2\theta)$, with a radial triangular kernel of radius 35 pixels, weighting each orientation by its coherency. The averaged vector $\overline{\mathbf{q}}$ is then normalized to unit length: $\widehat{\mathbf{q}} = \frac{\overline{\mathbf{q}}}{|\overline{\mathbf{q}}|}$. In each frame, we exclude positions with insufficient local orientation data or $|\overline{\mathbf{q}}| < 0.05$. We compare the remaining orientations at the same image positions: $C_{\mathrm{orientations}}(t) = \langle \widehat{\mathbf{q}}(\mathbf{x}, t_0) \cdot \widehat{\mathbf{q}}(\mathbf{x}, t) \rangle_{\mathrm{valid}}$. Here, $\langle \cdot \rangle_{\mathrm{valid}}$ denotes averaging over positions retained in both the reference and current frames. A decrease in $C_{\mathrm{orientations}}$ indicates departure from the reference pattern, while a subsequent increase indicates recovery toward it.

We check trigger events against the time-lapse images and estimate recovery times only for events whose correlation traces show a discernible post-wave rise and plateau. For each retained event, we define the start of recovery, $t_{\mathrm{min}}$, as the post-wave minimum of the smoothed correlation trace. We estimate the recovery endpoint, $t_{\mathrm{end}}$, from the intersection of two linear fits: one to the rising part of the trace and one to the subsequent plateau. The recovery time is $\tau_{\mathrm{recovery}} = t_{\mathrm{end}} - t_{\mathrm{min}}$.

### Local angular-relaxation model for repeated-wave nematic recovery

To interpret the approximately constant recovery times across repeated waves (Fig. 4e), we adapt the local angular dynamics of a cellular nematic coupled to a remodelable matrix[13]. Following the ordered, long-wavelength approximation of Ref.[13], we neglect spatial gradients and treat the densities and nematic-order magnitudes as constant during local post-wave recovery. Let $\phi_c(t)$ and $\phi_m(t)$ denote the cellular and mucilage-matrix director angles, with mismatch $\delta = \phi_c - \phi_m$. Their local dynamics are

$$2\frac{\mathrm{d}\phi_c}{\mathrm{d}t} = -\lambda_{\mathrm{pin}} \sin(2\delta), \qquad 2\frac{\mathrm{d}\phi_m}{\mathrm{d}t} = \lambda_{\mathrm{remodel}} \sin(2\delta). \tag{S23}$$

The pinning rate $\lambda_{\mathrm{pin}}$ describes cellular realignment toward the matrix, while $\lambda_{\mathrm{remodel}}$ describes reorientation of the matrix toward the cells. The corresponding ordered-state rate expressions are[13]

$$\lambda_{\mathrm{pin}} = \frac{k\beta_m \rho_m}{\beta_m \rho_m + \beta_c \rho_c}, \qquad \lambda_{\mathrm{remodel}} = k_- \rho_c. \tag{S24}$$

Here, $\beta_c$ and $\beta_m$ are the cell–cell and cell–matrix alignment strengths, $\rho_c$ and $\rho_m$ are the corresponding densities, $k$ is the cellular reorientation rate, and $k_-$ is the coefficient of cell-mediated matrix degradation in the original model. There, degradation and cell-oriented replacement allow the matrix to reorient. For diatoms, we interpret $\lambda_{\mathrm{remodel}}$ as an effective mucilage-rewriting rate.

Subtracting the angular equations gives

$$2\frac{\mathrm{d}\delta}{\mathrm{d}t} = -(\lambda_{\mathrm{pin}} + \lambda_{\mathrm{remodel}}) \sin(2\delta). \tag{S25}$$

For time-independent $\lambda_{\mathrm{pin}}$ and $\lambda_{\mathrm{remodel}}$ and nonperpendicular initial directors, the solution is

$$\tan\delta(t) = \tan\delta(t_{\mathrm{i}}) \exp\left[-\frac{t - t_{\mathrm{i}}}{\tau_\delta}\right], \qquad \tau_\delta = (\lambda_{\mathrm{pin}} + \lambda_{\mathrm{remodel}})^{-1}, \tag{S26}$$

where $t_{\mathrm{i}}$ marks the start of local recovery. We assume that cellular realignment is faster than mucilage rewriting during post-wave recovery. In the matrix-pinned limit, $\lambda_{\mathrm{pin}} \gg \lambda_{\mathrm{remodel}}$, the relaxation time is approximately

$$\tau_\delta \simeq \lambda_{\mathrm{pin}}^{-1}. \tag{S27}$$

To relate $\tau_\delta$ to the experimentally measured recovery time, we consider relaxation from mismatch $\delta_a$ to a smaller nonzero mismatch $\delta_b$. The required time is

$$\Delta t = \tau_\delta \ln\left|\frac{\tan\delta_a}{\tan\delta_b}\right|. \tag{S28}$$

The theoretical relaxation time thus provides a characteristic scale for the measured time:

$$\tau_{\text{recovery}} \sim \tau_\delta \simeq \lambda_{\text{pin}}^{-1}, \tag{S29}$$

Therefore, the recovery times remain consistent with an unchanged post-wave pinning rate.

### Numerical simulations of trigger-wave-induced nematic memory remodeling

We use a two-dimensional phenomenological model to examine trigger-wave propagation, cellular realignment and remodeling of the mucilage matrix. The model couples activator–recovery kinetics to nematic-guided motion, cue-dependent weakening of matrix pinning and slow reorientation of the matrix.

The simulated fields are the cell density $\varrho(\mathbf{x}, t)$, activator $a(\mathbf{x}, t)$, recovery variable $b(\mathbf{x}, t)$, activation cue concentration $c(\mathbf{x}, t)$, phenomenological scalar cellular alignment amplitude $S_{\text{cell}}(\mathbf{x}, t)$, cellular director angle $\varphi(\mathbf{x}, t)$, and mucilage-matrix director angle $\varphi_m(\mathbf{x}, t)$, where $\mathbf{x} = (x, y)$ denotes position in the simulation plane. The cellular and matrix directors are $\mathbf{n}_{\text{cell}} = (\cos\varphi, \sin\varphi)$, and $\mathbf{n}_{\text{mat}} = (\cos\varphi_m, \sin\varphi_m)$. Both fields are nematic, so $\varphi \equiv \varphi + \pi$ and $\varphi_m \equiv \varphi_m + \pi$. The field $\varphi_m$ represents the orientational memory stored in the mucilage matrix. Unlike the cellular director, it is not advected by cell motion.

The initial matrix texture is prescribed as an isolated nematic defect, $\varphi_m(\mathbf{x}, 0) = m_{\text{d}}\vartheta(\mathbf{x}) + \varphi_0$, where $m_{\text{d}}$ is the defect charge, $(x_c, y_c)$ is the defect center, $\vartheta(\mathbf{x}) = \text{atan2}(y - y_c, x - x_c)$ is the polar angle around the defect, and $\varphi_0$ sets the defect orientation. The cellular director is initially aligned with the matrix, $\varphi(\mathbf{x}, 0) = \varphi_m(\mathbf{x}, 0)$. The memory-remodeling simulations use an initial $m_{\text{d}} = -1/2$ texture, whereas the defect-focused simulations use both $m_{\text{d}} = +1/2$ and $m_{\text{d}} = -1/2$ textures. A trigger wave is initiated by imposing a small localized region of elevated activator $a$ and cue concentration $c$ near one side of the domain. For the defect-focused simulations, the defect orientation, $\varphi_0$, and trigger position are chosen to reproduce the relative front–defect geometries shown in Fig. 3f,g. In the $+1/2$ case, the wave approaches the converging side of the polar texture, showing the focusing mode. We note that this response is incidence-dependent: waves approaching from the opposite or sufficiently off-axis side may be redirected around the core. Such trajectories remain defect-guided because the surrounding director field can shape the wave without requiring passage through the singular core.

**Excitable trigger wave.** We use FitzHugh–Nagumo-type activator–recovery kinetics[14, 15], with additional feedback from the activation cue. The activator $a$ and recovery variable $b$ are defined per cell. We write their transport and reaction equations in terms of the density-weighted fields $\varrho a$ and $\varrho b$:

$$\partial_t \varrho + \nabla \cdot (\varrho \mathbf{u}) = D_\varrho \nabla^2 \varrho, \tag{S30}$$

$$\partial_t(\varrho a) + \nabla \cdot (\varrho a \mathbf{u}) = D_a \nabla^2 (\varrho a) + \varrho \frac{a(1-a)(a-a_*) - b + \lambda_{ac} c}{\tau_a}, \tag{S31}$$

$$\partial_t(\varrho b) + \nabla \cdot (\varrho b \mathbf{u}) = D_b \nabla^2 (\varrho b) + \varrho \epsilon_b (a - r_b b). \tag{S32}$$

Here $\mathbf{u}(\mathbf{x}, t)$ is the coarse-grained cellular velocity. The coefficients $D_\varrho$, $D_a$ and $D_b$ govern diffusion of $\varrho$, $\varrho a$ and $\varrho b$, respectively. $a_*$ sets the cubic activator threshold, $\lambda_{ac}$ controls feedback from the activation cue to the activator, $\tau_a$ is the activator timescale, $\epsilon_b$ is the recovery rate, and $r_b$ sets the relaxation strength of the recovery variable.

We model the activation cue concentration $c(\mathbf{x}, t)$ through production by activated cells, diffusion and decay, assuming that the cue is advected at the cellular velocity $\mathbf{u}$:

$$\partial_t c + \nabla \cdot (c \mathbf{u}) = D_c \nabla^2 c + k_c \varrho H(a - a_{\mathrm{th}}) - \frac{c}{\tau_c}. \tag{S33}$$

In this equation, $D_c$ is the cue diffusivity, $k_c$ is the cue-production rate, $H$ is a Heaviside threshold function, $a_{\mathrm{th}}$ is the activator threshold for cue production, and $\tau_c$ is the cue-decay timescale.

**Cue-dependent detachment and nematic-guided motion.** The cue concentration controls the local attachment state between cells and the mucilage matrix. We introduce a detachment variable

$$g_{\mathrm{detach}}(c) = \frac{c}{c + K_c}, \tag{S34}$$

where $K_c$ is the cue half-saturation constant. Thus, $g_{\mathrm{detach}}(c) \approx 1$ represents a detached, motile, matrix-writing state. By contrast, $1 - g_{\mathrm{detach}}(c)$ represents an attached, matrix-pinned state.

We constrain population motion to the local nematic axis and choose its direction toward lower cue concentration. The polarity rule used in the simulations is

$$\zeta = -\tanh\left(\frac{\mathbf{n}_{\mathrm{cell}} \cdot \nabla c}{g_c}\right), \tag{S35}$$

where $g_c$ sets the cue-gradient scale over which polarity reverses. The resulting coarse-grained cellular velocity is

$$\mathbf{u} = U(c)\, S_{\mathrm{cell}}\, \zeta\, \mathbf{n}_{\mathrm{cell}}, \qquad U(c) = U_0 + U_1 g_{\mathrm{detach}}(c). \tag{S36}$$

Here $U(c)$ is the cue-dependent motility scale, $U_0$ is the basal motility coefficient, and $U_1$ is the cue-enhanced motility coefficient. The factor $S_{\rm cell}$ acts as a nematic coherence factor: highly ordered regions generate coherent advective transport along the local director, whereas weakly ordered regions generate little net coarse-grained flux even if individual cells remain motile.

**Strength of local cellular alignment.** We represent local cellular nematic order by a phenomenological scalar amplitude $S_{\rm cell}$, while $\varphi$ specifies the director angle. We use a nonconserved relaxation law[16], with $S_{\rm cell}$ relaxing toward a preferred local value $S_{\rm tar}$:

$$\partial_t S_{\rm cell} = \frac{S_{\rm tar} - S_{\rm cell}}{\tau_S} + D_S \nabla^2 S_{\rm cell}, \tag{S37}$$

where $\tau_S$ is the local relaxation timescale and $D_S$ controls spatial coupling of the scalar amplitude. We choose $S_{\rm tar}$ to favor stronger alignment in dense, matrix-attached regions and weaker alignment during activation or strong director distortion. At each position and time, we define $S_{\rm tar}$ through a phenomenological balance between recovery toward a baseline amplitude $S_{\rm base}$ and loss of alignment due to the cue and director distortion:

$$r_{\rm rec}(S_{\rm base} - S_{\rm tar}) = \left[r_c c + r_\nabla |\nabla_N \varphi|^2\right] S_{\rm tar}. \tag{S38}$$

Here, $r_{\rm rec}$ is the recovery rate toward $S_{\rm base}$. The coefficients $r_c$ and $r_\nabla$ control cue- and distortion-dependent disordering, respectively. The operators $\nabla_N$ and $\nabla_N^2$ denote angular gradients and Laplacians evaluated through the doubled-angle components $\cos 2\varphi$ and $\sin 2\varphi$, with the same construction for $\varphi_m$. The baseline amplitude is $S_{\rm base} = S_0(\varrho/\varrho_0)^{\gamma_\varrho} G_{\rm pin}(c)$, where $G_{\rm pin}(c) = 1 - g_{\rm detach}(c)$. Here, $S_0$ is the reference amplitude, $\varrho_0$ is the reference density and $\gamma_\varrho$ controls the density dependence. Solving the local balance gives

$$S_{\rm tar} = \frac{S_0(\varrho/\varrho_0)^{\gamma_\varrho} G_{\rm pin}(c)}{1 + \alpha_S c + \kappa_S |\nabla_N \varphi|^2}, \tag{S39}$$

where $\alpha_S = r_c/r_{\rm rec}$ and $\kappa_S = r_\nabla/r_{\rm rec}$.

**Dynamics of cellular director and hidden mucilage-memory.** We represent cellular realignment and matrix remodeling by coupled nematic torques, following the cell–matrix framework in Ref.[13]. The cue-dependent factors are phenomenological additions. The cellular director evolves through advection, angular smoothing, alignment toward the matrix and rotational fluctuations:

$$\partial_t \varphi + \mathbf{u} \cdot \nabla_N \varphi = D_\varphi \nabla_N^2 \varphi - \frac{1 - g_{\rm detach}(c)}{\tau_{\rm align}[1 + \beta_{\rm act}\mathcal{A}]} \sin[2(\varphi - \varphi_m)] + \xi_\varphi. \tag{S40}$$

Here $D_\varphi$ is the cellular angular diffusivity and $\tau_{\rm align}$ sets the strength of cellular alignment toward the matrix. We define the motility-dependent activity as $\mathcal{A} = [U(c)|\zeta|]^2$ during the wave-active period and set $\mathcal{A} = 0$ when cellular motility is switched off. The anchoring torque, proportional to $-\sin[2(\varphi - \varphi_m)]$, rotates the cellular director toward the local matrix director. Rising cue concentration weakens this torque through $1 - g_{\mathrm{detach}}(c)$, while $\beta_{\mathrm{act}}$ controls its further suppression by activity.

The term $\xi_\varphi$ represents rotational noise in the cellular director angle. We assume that motility enhances these fluctuations and that increasing cell density reduces their activity-dependent contribution. Where $S_{\mathrm{cell}} \leq 0.4$, the cellular director evolves according to Eq. (S40). We implement this noise term by adding independent, zero-mean Gaussian angular increments at each grid point and time step, with variance $2D_{\mathrm{rot}}\Delta t$, where $D_{\mathrm{rot}} = D_{\mathrm{rot},0} + D_{\mathrm{rot},1}\frac{\mathcal{A}}{\mathcal{A}+\mathcal{A}_0}\frac{\varrho_{\mathrm{ref}}}{\varrho+\varrho_\epsilon}$. Here, $D_{\mathrm{rot},0}$ and $D_{\mathrm{rot},1}$ set the basal and activity-dependent noise strengths, respectively. The parameters $\mathcal{A}_0$ and $\varrho_{\mathrm{ref}} = \varrho_0$ are the activity saturation scale and reference density, while the positive offset $\varrho_\epsilon$ keeps the density factor finite at low density. Where $S_{\mathrm{cell}} > 0.4$, we instead enforce alignment with the matrix by setting $\varphi = \varphi_m$ after the cellular-angle update and before updating the matrix. This constraint overrides both the deterministic rotation and the random increment, so the matrix-rewriting torque vanishes at these positions during that update.

The mucilage matrix is modeled as a hidden, non-advected nematic memory field. Its director evolves by passive angular smoothing and by cue-gated rewriting toward the instantaneous cellular director:

$$\partial_t \varphi_m = D_{\mathrm{mat}} \nabla_N^2 \varphi_m + \frac{g_{\mathrm{detach}}(c)}{\tau_{\mathrm{write}}} \sin[2(\varphi - \varphi_m)]. \tag{S41}$$

Here, $D_{\mathrm{mat}}$ controls the passive relaxation of spatial variations in the matrix director angle, and $1/\tau_{\mathrm{write}}$ sets the strength of cue-dependent rewriting. The rewriting term rotates the matrix toward the local cellular director when the two are misaligned and the cue is present. As the cue decays, rewriting weakens and cells realign toward the current matrix orientation.

**Numerical implementation and simulation protocols.** The equations are solved on a square domain of size $L_x = L_y = 12$ using a $300 \times 300$ grid and an explicit time step $\Delta t = 2 \times 10^{-3}$. Advective terms in $\varrho$, $\varrho a$, $\varrho b$, and $c$ are discretized using a conservative upwind finite-volume scheme, and diffusion terms are evaluated using a five-point Laplacian with zero-gradient boundary conditions. During integration, we impose the numerical bounds $\varrho \geq 10^{-4}$, $0 \leq a \leq 1.5$, $b, c \geq 0$ and $0 \leq S_{\mathrm{cell}} \leq 1.2$. The density floor prevents division by zero when calculating the per-cell variables. The remaining bounds enforce nonnegative states and cap the activator and alignment amplitudes.

The memory-remodeling simulation starts from an $m_{\mathrm{d}} = -1/2$ texture with $D_{\mathrm{mat}} = 0$ and $\tau_{\mathrm{write}} = 10$. The matrix therefore changes only through cue-dependent rewriting, without passive angular smoothing. To examine post-wave recovery, we switch off new cue production and cellular motility at $t = 35$ and increase $D_\varrho$ from $10^{-4}$ to $5 \times 10^{-2}$ to accelerate density relaxation. The existing cue continues to diffuse and decay, while the remaining fields evolve during this imposed recovery stage. Separate defect-focused simulations use $m_\mathrm{d} = +1/2$ and $m_\mathrm{d} = -1/2$ textures to visualize how trigger waves propagate around nematic defects and where the cellular director becomes disrupted. Complete parameter values are provided in the accompanying simulation scripts.

Recovery and remodeling are quantified using nematic orientational correlations relative to the initial state: $C_\mathrm{m}(t) = \langle \cos\{2[\varphi_m(\mathbf{x},t) - \varphi_m(\mathbf{x},0)]\}\rangle_\mathbf{x}$, and $C_\mathrm{cells}(t) = \langle \cos\{2[\varphi(\mathbf{x},t) - \varphi(\mathbf{x},0)]\}\rangle_\mathbf{x}$. Here, $\langle\cdot\rangle_\mathbf{x}$ denotes a spatial average over the simulation domain. Values near one indicate retention or recovery of the initial texture; lower values indicate a departure from it.

### Raphe morphology and extracellular material in *Cylindrotheca* sp.

Extracellular polymers secreted through the raphe support substrate adhesion and traction during pennate-diatom gliding[17, 18]. Scanning electron microscopy (SEM) of our *Cylindrotheca* sp. isolate reveals abundant extracellular material coating cell surfaces and bridging neighboring cells (Fig. S7a), together with a longitudinal slit in the silica frustule consistent with the raphe (Fig. S7b). The extracellular material is morphologically consistent with mucilage, although SEM alone does not establish its chemical composition. Wheat germ agglutinin (WGA) staining provides complementary evidence for a glycoconjugate-rich extracellular matrix (Fig. 4d).

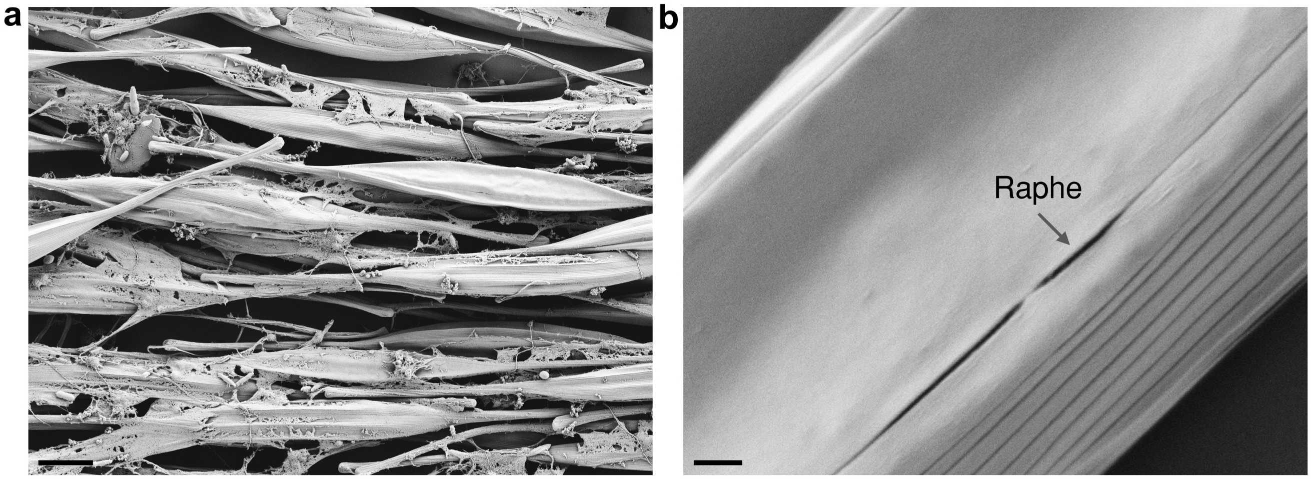


**Fig. S7. The raphe and extracellular material of *Cylindrotheca* sp.** (**a**) Scanning electron micrograph showing abundant extracellular material coating and bridging densely packed cells. (**b**) Higher-magnification view of the longitudinal raphe slit in the silica frustule (arrow). Scale bars, $4\ \mu\mathrm{m}$ (**a**) and $300\ \mathrm{nm}$ (**b**).

#### Nematic memory preserves large-scale architecture while allowing local reorientation

To assess how closely the director pattern recovers between waves, we compare the same field immediately before two recurrent trigger waves separated by approximately $8\ \mathrm{h}$ (Fig. S8). The large-scale director architecture remains recognizable, with modest orientation changes across most of the field. Larger shifts are concentrated in a few defect-associated regions, where some defect patterns become less distinct (Figs. 4g and S8).

Together with the persistence of oriented mucilage deposited by the cells (Fig. 4d), these observations support a material-memory mechanism in which the extracellular matrix preserves a broad orientational template, while recurrent waves locally remodel the cellular nematic architecture.

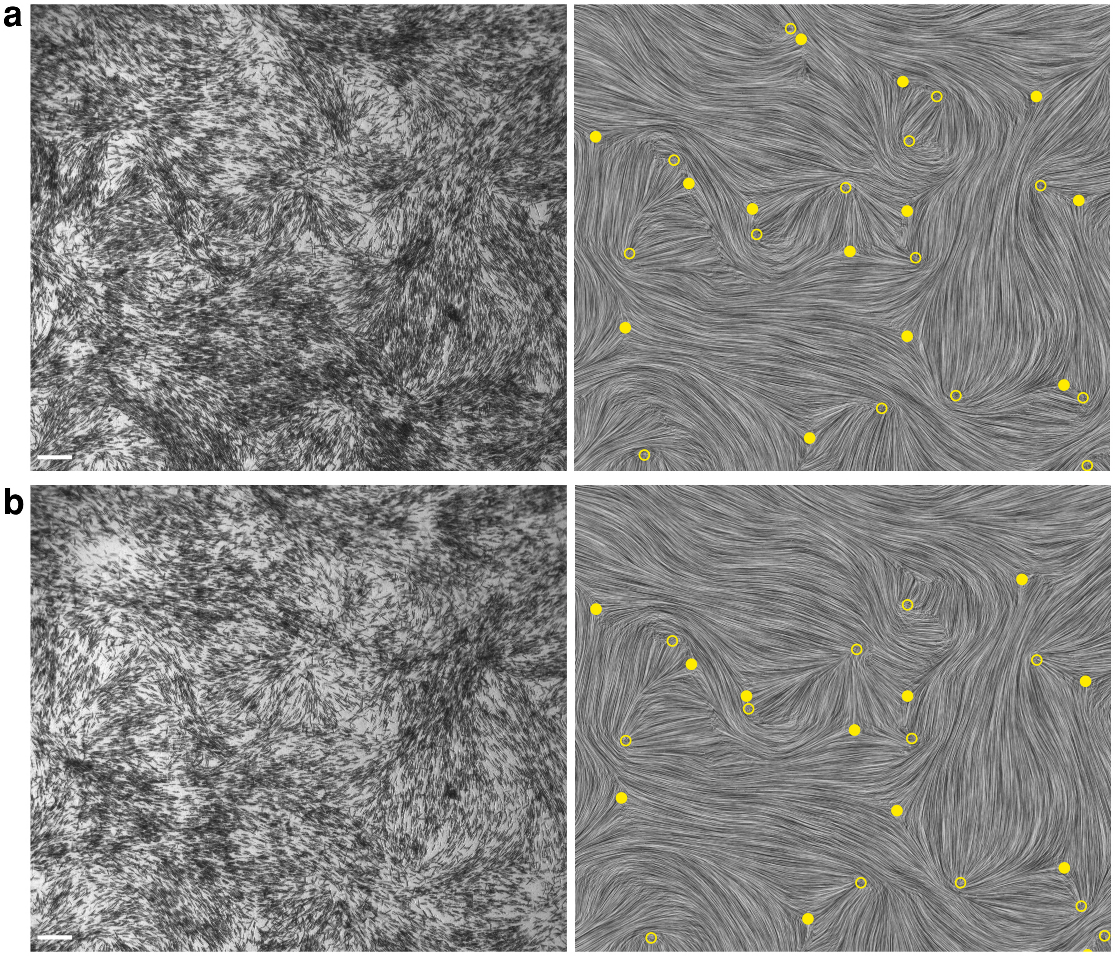


**Fig. S8. Large-scale nematic architecture persists despite local reorientation.** (**a,b**) The same field of view immediately before the first (**a**) and second (**b**) recurrent trigger waves, separated by approximately $8\ \mathrm{h}$. Left, bright-field images; right, coarse-grained director textures rendered by line-integral convolution. Open and filled yellow symbols denote $+1/2$ and $-1/2$ defects, respectively. The broad director architecture remains recognizable, with local changes in defect-associated patterns. Scale bars, $100\ \mu\mathrm{m}$.

### Trigger waves displace and progressively remodel topological defects

Topological defects shape trigger-wave pathways. To examine how wave passage modifies this landscape, we track an individual defect during one wave encounter and quantify changes in defect positions and counts across recurrent waves.

In the representative encounter shown in Fig. S9a, manual tracking shows that the advancing wave displaces a $+1/2$ defect by approximately 140 $\mu\mathrm{m}$ (Fig. S9b). The defect subsequently moves back toward its pre-wave position but does not fully return during the observation period.

To quantify cumulative remodeling, we compare the defects detected in each frame with those in the first analyzed frame, at time $t_0$. We pair defects of the same charge using one-to-one nearest-neighbor matching within 126 $\mu\mathrm{m}$. This cutoff is comparable to the orientational correlation length $\xi$ and smaller than the defect-density length $\ell_{\mathrm{defect}}$. We classify paired defects as matched, unmatched current defects as emerged and unmatched reference defects as disappeared. These categories describe matches to the reference frame and do not by themselves establish defect creation or annihilation.

For matched defects, the mean displacement from their initial positions is

$$\bar{d}_{\mathrm{def}}(t) = \frac{1}{N_{\mathrm{match}}(t)} \sum_{i=1}^{N_{\mathrm{match}}(t)} \|\mathbf{r}_i(t) - \mathbf{r}_i(t_0)\| \,, \tag{S42}$$

where $\mathbf{r}_i$ is the defect position and $N_{\mathrm{match}}(t)$ is the number of matches to the reference at time $t$. The set of defects contributing to this average can therefore change over time.

Across recurrent waves, the mean displacement increases, with partial decreases after individual events (Fig. S9c). The number of matched defects decreases and the number classified as disappeared increases, while emerged defects remain comparatively sparse. The total number of detected defects also declines (Fig. 4h).

To map director reorientation that persists between waves, we compare the two relaxed pre-wave states shown in Fig. 4g. At each position, we calculate the axial angular mismatch:

$$\Delta\theta(\mathbf{x}) = \arccos|\cos\left[\theta_2(\mathbf{x}) - \theta_1(\mathbf{x})\right]| \,, \qquad 0 \leq \Delta\theta \leq \frac{\pi}{2}, \tag{S43}$$

where $\theta_1$ and $\theta_2$ are the local director angles in the two states. Only positions with a defined director angle in both states are included.

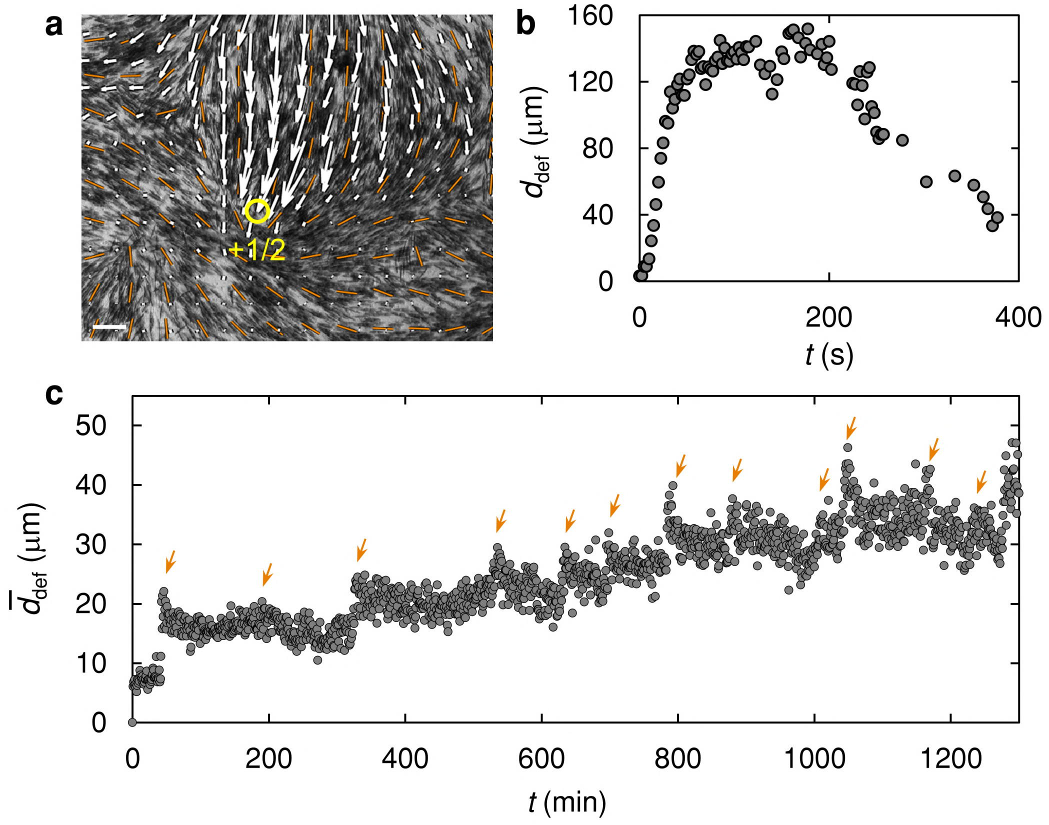


**Fig. S9. Trigger waves displace nematic defects and progressively remodel the topological landscape.** (**a**) Representative encounter between a propagating trigger wave and a $+1/2$ defect. The yellow circle marks the defect core, orange segments show the local nematic director, and white arrows show the wave-associated PIV velocity field. Scale bar, $50\ \mu\mathrm{m}$. (**b**) Displacement, $d_{\rm def}(t)$, of the marked defect core from its pre-wave position during the event in (**a**), measured by manual tracking. The defect reaches a displacement of approximately $140\ \mu\mathrm{m}$ and subsequently returns only partway toward its initial position. (**c**) Mean displacement, $\overline{d}_{\rm def}(t)$, of same-charge defects matched to the first analyzed frame. Orange arrows mark trigger waves. The mean displacement increases overall, with transient decreases after some waves.

### Topographic control of diatom alignment

To test whether substrate topography can prescribe the cellular director field, we fabricate patterned polydimethylsiloxane (PDMS) surfaces using maskless photolithography and replica molding. The grooves are 11–12 $\mu\mathrm{m}$ deep and include smoothly curved, sinusoidal and character-like patterns (Fig. S10).

*Cylindrotheca* sp. cells align tangentially to the grooves and follow their changing curvature, allowing the substrate geometry to guide the cellular director field (Figs. 5b and S10a). On the "Stanford" pattern, cells remain close to the letter-shaped groove and align along its contour at low density (Figs. 5b and S10b). As the population grows, the occupied band broadens beyond the groove and the imposed orientation pattern becomes less distinct (Fig. S10b). By contrast, the prescribed alignment persists at higher density in arrays of closely spaced grooves.

These observations suggest a competition between local groove confinement and collective nematic alignment. As the populated region expands beyond an isolated groove, topographic guidance becomes less effective, whereas closely spaced grooves maintain confinement across the mat. Pattern fidelity therefore appears to depend on groove spacing relative to the lateral extent of collective nematic organization.

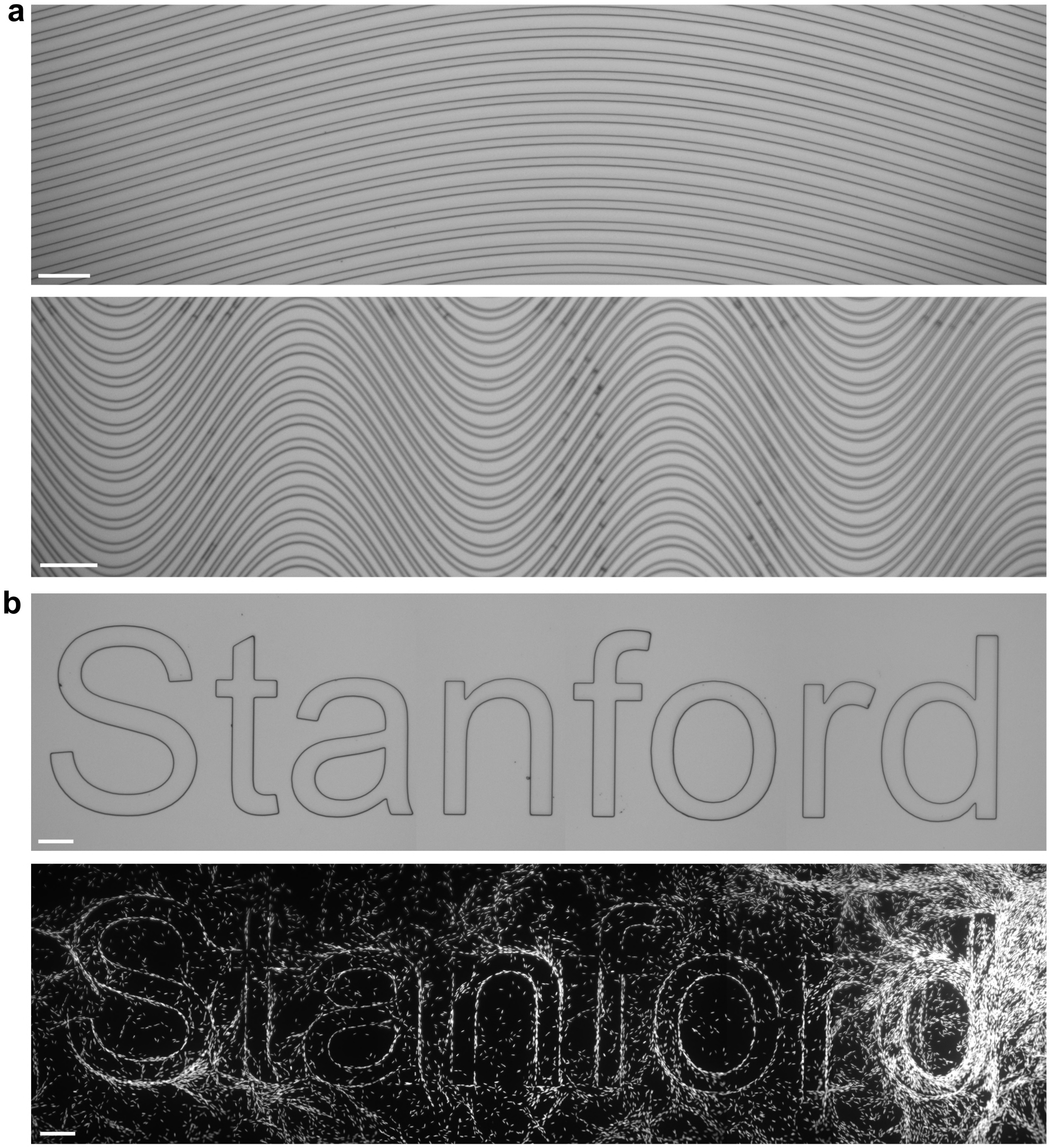


**Fig. S10. Topographic control of diatom alignment.** (**a**) Representative curved (top) and sinusoidal (bottom) PDMS microgroove patterns used to impose orientation fields with varying local curvature. Scale bars, 100 $\mu$m. (**b**) PDMS topography spelling "Stanford" before cell seeding (top) and the corresponding organization of *Cylindrotheca* sp. after colonization (bottom). Cells follow the letter contours most clearly in lower-density regions, whereas the cellular pattern becomes less distinct at higher density. Scale bars, 200 $\mu$m.

**Table S1.** Exponential fits to the orientational correlation function. Fits use $C_{\mathrm{conn}} = A_0 \exp(-r/\xi)$. Fields for which the fitted separation is less than $2\xi$ are identified as field-of-view (FOV) limited. The eight independent datasets contribute to the reported image-level mean $\xi = 123.33 \pm 7.98\ \mu\mathrm{m}$.

| Dataset | $\xi$ ($\mu$m) | $A_0$ | $R^2$ | Fitted range ($\mu$m) | $N_{\mathrm{bin}}$ | FOV status |
|---|---|---|---|---|---|---|
| 20× | 108.12 | 0.93 | 0.9878 | 182.63 | 31 | Limited |
| 20× | 122.11 | 0.87 | 0.9996 | 236.06 | 40 | Limited |
| 20× | 154.44 | 0.84 | 0.9990 | 182.63 | 31 | Limited |
| Stitched 20× | 102.02 | 0.89 | 0.9929 | 1188.41 | 199 | OK |
| 4× | 146.08 | 0.89 | 0.9886 | 925.13 | 155 | OK |
| 4× | 131.45 | 0.65 | 0.9348 | 925.13 | 155 | OK |
| 4× | 88.47 | 1.01 | 0.9897 | 925.13 | 155 | OK |
| 4× | 133.94 | 0.91 | 0.9910 | 1189.31 | 199 | OK |

**Supplementary Movie Legends**

**Supplementary Movie 1** Tidal-flat diatoms form nematic mosaics that support trigger waves

**Supplementary Movie 2** Single-cell tracking reveals two motility states

**Supplementary Movie 3** Director-guided, density-gated trigger-wave routing

**Supplementary Movie 4** Topological defect landscape guides and is remodeled by trigger waves

**Supplementary Movie 5** Recurrent trigger waves with partial recovery of nematic alignment

**Supplementary Movie 6** Diatom-deposited mucilage retains a hidden orientational template

**Supplementary Movie 7** Mechanical, chemical and optical perturbations nucleate trigger waves
